# Interplay of folded domains and the disordered low-complexity domain in mediating hnRNPA1 phase separation

**DOI:** 10.1101/2020.05.15.096966

**Authors:** Erik W. Martin, F. Emil Thomasen, Nicole M. Milkovic, Matthew J. Cuneo, Christy R. Grace, Amanda Nourse, Kresten Lindorff-Larsen, Tanja Mittag

## Abstract

Liquid-liquid phase separation underlies the membrane-less compartmentalization of cells. Intrinsically disordered low-complexity domains (LCDs) often mediate phase separation, but how their phase behavior is modulated by folded domains is incompletely understood. Here, we interrogate the interplay between folded and disordered domains of the RNA-binding protein hnRNPA1. The LCD of hnRNPA1 is sufficient for mediating phase separation *in vitro*. However, we show that the folded RRM domains and a folded solubility-tag modify the phase behavior, even in the absence of RNA. Notably, the presence of the folded domains reverses the salt dependence of the driving force for phase separation relative to the LCD alone. Small-angle X-ray scattering experiments and coarse-grained MD simulations show that the LCD interacts transiently with the RRMs and/or the solubility-tag in a salt-sensitive manner, providing a mechanistic explanation for the observed salt-dependent phase separation. These data point to two effects from the folded domains: (1) electrostatically mediated interactions that compact hnRNPA1 and contribute to phase separation, and (2) increased solubility at higher ionic strengths mediated by the folded domains. The interplay between disordered and folded domains can modify the dependence of phase behavior on solution conditions and can obscure signatures of physicochemical interactions underlying phase separation.

**Graphical abstract:** hnRNPA1 phase separation is highly salt sensitive.
Phase separation of the low-complexity domain (LCD) of hnRNPA1 increases with NaCl. In contrast, phase separation of full-length hnRNPA1 is salt-sensitive. At low NaCl concentrations, electrostatic RRM-LCD interactions occur and can contribute positively to phase separation, but they are screened at high NaCl concentrations. The folded domains solubilize hnRNPA1 under these conditions and prevent phase separation.

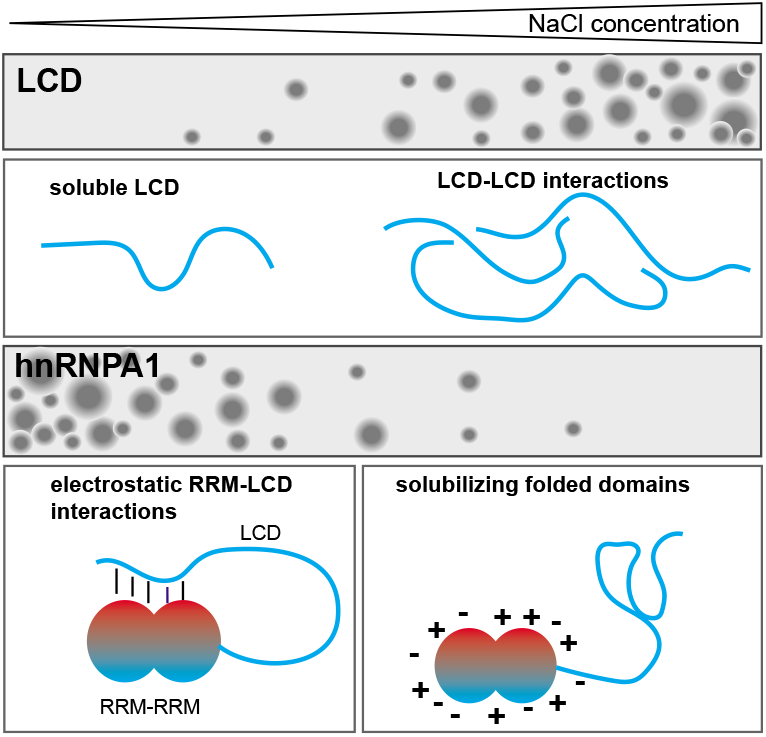

## Introduction

Liquid-liquid phase separation (LLPS) mediates the extensive compartmentalization of cells and leads to the formation of membraneless organelles including nucleoli, stress granules and P bodies amongst many others (1–3). In addition to typical membraneless organelles, other biomolecular condensates that are formed via LLPS include heterochromatin (4–6), transcriptional condensates (7–9) and membrane receptor clusters (10,11). Understanding the interactions underlying phase separation therefore has the potential to provide insights into a wide variety of fundamental biological processes. Associative phase separation is driven by multivalent interactions such as those that occur between tandem repeats of folded domains and linear motifs in pairs of proteins, or between short motifs in intrinsically disordered low-complexity domains (LCDs) (12–17). These multivalent interactions mediate the formation of three-dimensional protein networks whose formation is typically coupled to a density transition that results in dilute and dense coexisting phases (12,18).

Recent progress has improved our understanding of how phase behavior is encoded in LCDs of RNA-binding proteins (13,15,17,19,20), which comprise a high fraction of small polar residues and are interspersed with aromatic and few charged residues. We showed that aromatic residues contribute the main driving force for phase separation in prion-like LCDs (17). However, LCDs rarely exist in isolation and are typically connected to folded domains, either as tails or internal linkers between folded domains. How such architectures modulate the phase behavior of LCDs remains largely unexplored although it is exploited experimentally. For example, soluble folded domains can increase the saturation concentration of LCDs and can thus be used to prevent phase separation until the solubilizing domain is proteolytically cleaved (21–23). However, it is often assumed that the direct fusion of an LCD with a fluorescent protein or reporter domain does not dramatically influence its phase behavior (7,24). Here, we directly address this question by determining the salt dependent phase behavior of the LCD of hnRNPA1 in isolation and in the context of the full-length protein and with the addition of a solubility tag.

hnRNPA1, an archetypal member of the heterogeneous nuclear ribonuclear protein (hnRNP) family, which shuttles in and out of the nucleus, associates with pre-mRNA and acts as a splicing factor (25). Under stress conditions, hnRNPA1 is sequestered in cytoplasmic stress granules, which are formed via liquidliquid phase separation (26). Mutations in the hnRNPA1 LCD lead to familial forms of amyotrophic lateral sclerosis and multisystem proteinopathy (27), two related diseases that are characterized by solid deposits of RNA-binding proteins and lead to neurodegeneration. hnRNPA1 comprises tandem RNA recognition motifs (RRMs) that behave like a single folded module due to their short connecting linker (28) and a long, intrinsically disordered LCD. This domain architecture is typical for many RNA-binding proteins, and hnRNPA1 thus serves as an archetypal member of a large family of proteins.

hnRNPA1 undergoes phase separation with RNA via multivalent interactions mediated by its two RRM domains and RGG motifs in the LCD. In the absence of RNA, the LCD is necessary and sufficient for phase separation (26). Here, we characterize the salt-dependence of phase separation of full-length hnRNPA1, domain deletion constructs and constructs with an additional folded domain from a solubility tag to characterize the nature of the interactions that drive phase separation. Our results show that the solution conditions enabling LCD phase separation are highly susceptible to the presence of folded domains, likely because the solubility profiles of folded domains and LCDs as a function of solution conditions differ strongly and because of interactions between the LCD and folded domains. Small-angle X-ray scattering (SAXS) measurements demonstrate a small increase of the dimensions of hnRNPA1 at salt concentrations at which electrostatic interactions are shielded, and coarse-grained molecular dynamics (MD) simulations provide direct evidence for transient interactions between the LCD and the folded domains. These interactions modulate the solution conditions under which the protein can undergo phase separation. As a result, the salt dependence of hnRNPA1 phase behavior is not directly diagnostic of the interactions mediating phase separation, but also depends on the effects of the folded domains. Given that folded domains have co-evolved with intrinsically disordered domains in many proteins, they may encode functional interactions between the two (29) that help regulate phase separation.

## MATERIAL AND METHODS

### Protein Expression and Purification

hnRNPA1* (where * denotes that the hexa-peptide 259-264 is deleted) protein and deletion constructs were expressed as N-terminally tagged hSUMO fusion proteins in BL21 (DE3) RIPL cells (Agilent) in LB media, and purified using Ni^2+^ affinity chromatography, followed by size exclusion chromatography (SEC), RNaseA digestion followed immediately by ion exchange, and SEC, as previously described (26). Briefly, cells were lysed in lysis/wash buffer (50 mM HEPES pH 7.5, 250 mM NaCl, 30 mM imidazole, 2 mM 2-mercaptoethanol (βME), and complete protease inhibitor cocktail) (Roche) with a microfluidizer (Microfluidics, 20,000 psi). Clarified lysate was filtered and passed over a Ni^2+^ affinity chromatography column by gravity and washed with the lysis/wash buffer. Additional wash steps containing 5 column volumes of wash buffer with 1 mg/mL RNaseA followed by 5 column volumes of wash buffer containing 600 mM NaCl were used to remove RNA. A final wash step containing 50 mM imidazole was included prior to eluting the protein with a buffer containing 50 mM HEPES pH 7.5, 300 mM NaCl, 300 mM imidazole, 2 mM βME. Protein-containing fractions were concentrated and passed over a size-exclusion chromatography (SEC) column (Superdex 200 16/60 column (hSUMO-hnRNPA1*) or a Superdex 75 (16/60) (hSUMO-LCD*)) equilibrated in storage buffer (50 mM HEPES pH 7.5, 300 mM NaCl, 5 mM DTT.) If residual contaminates remained, proteins were further purified by ion exchange chromatography on a 5mL HiTrap SP. The hSUMO tag was cleaved off the hSUMO-hnRNPA1* protein by treatment with Ulp1 at room temperature for 2 hours. Ulp1 and the hSUMO tag were removed by SEC.

The LCD* was expressed as His-tagged fusion protein and purified from inclusion bodies as previously reported (17). The His-tag was cleaved off by TEV cleavage and removed by Ni-NTA and SE chromatography.

All proteins were concentrated using ultrafiltration concentrators with either 3 kDa or 30 kDa MWCO regenerated cellulose membrane (EMD Millipore), flash frozen in small aliquots in liquid nitrogen, and stored at −80 °C until needed. Protein identity was determined by mass spectrometry, purity was determined by Coomassie Blue-stained SDS-PAGE gel and the monodispersity of samples was confirmed by dynamic light scattering. RNA content was analyzed by the ratio of the absorbance at 260 and 280 nm and/or polyacrylamide gel.

### Differential Interference Contrast (DIC) Microscopy

Protein samples were diluted to 300 μM in 50 mM HEPES pH 7.5, 5 mM DTT and varying salt concentrations (50, 100, 200, 300 mM NaCl). Sealed sample chambers containing protein solutions comprised a microscope slide and a coverslip, sandwiching 3M 300 LSE high-temperature double-sided tape (0.34 mm) with windows for microscopy cut out. Droplets were observed on a Nikon C2 laser scanning confocal microscope with a 20X (0.8NA) Plan Apo objective. Images were processed with the Nikon NIS Elements software. All images within the same row of figures were taken with the same camera settings.

### *In Vitro* Determination of Phase Diagrams

Dilute phase concentrations were determined as reported previously (30). All protein constructs were purified and stored in storage buffer as above. Protein was diluted with a buffer containing no salt (50 mM HEPES pH 7.5, 5 mM DTT) to induce LLPS. The samples were then passed through 0.22 μm filters (4 mm diameter) to remove any particulate matter that could nucleate LLPS or aggregate formation. The samples were partitioned into 12 μL aliquots into clear, colorless tubes and incubated at the desired temperatures for 20 min. The dense phases in the temperature-equilibrated samples were then sedimented in a temperature-equilibrated centrifuge for 5 min at maximum speed (21,000 g). 7 μL of the resulting supernatant (i.e. the dilute phase) was gently removed and placed into a clean tube. The supernatants were then diluted two-fold with an appropriately matched buffer to ensure that the sample does not undergo LLPS at room temperature. The protein concentration of the dilute phase was determined from the absorbance at 280 nm on a NanoDrop UV/Vis spectrophotometer. Phase diagrams were determined for the various protein constructs at 50, 75, 100, 125, 150 mM NaCl and 5, 10, 15, 20, 25, and 30 °C. Each coexistence curve was fitted to the scaling relation for binary demixing adapted from renormalization-group theory (31–33):

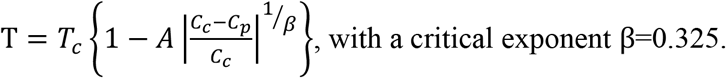

### Small Angle X-Ray Scattering (SAXS) Sample Preparation and Data Collection

Samples of full-length hSUMO-hnRNPA1* and hnRNPA1* were prepared in a buffer containing 50 mM Tris pH 7.5, 300 mM NaCl, 10 mM DTT, 2 mM TCEP. DTT and TCEP were used to scavenge radicals and prevent radiation damage. Samples of LCD* were prepared in 6M GndCl and exchanged into 50 mM Tris pH 7.5, 10 mM DTT, 2 mM TCEP, and a range of NaCl concentrations by SEC. SAXS data were collected as a function of NaCl concentration. Experiments were performed at the BioCAT (beamline 18ID at the Advanced Photon Source, Chicago) with in-line size exclusion chromatography (SEC-SAXS) to separate monomeric protein from aggregates and ensure the best possible buffer subtraction. Concentrated protein samples were injected into a Superdex 200 or Superdex 75 increase column (GE Lifesciences) preequilibrated in a buffer containing 50 mM Tris pH 7.5, 10 mM DTT, 2 mM TCEP, and the desired NaCl concentration, using an FPLC running at 0.4 mL/min. The output of the column passed through UV and conductance monitors before injection into a coflow sample chamber. The coflow sample chamber sheaths the sample in a jacket of matched buffer, preventing radiation damage (34,35). Scattering intensity was recorded using a Pilatus3 1M (Dectris) detector placed 3.5 m from the sample providing access to a q-range from 0.004-0.4 Å^-1^. One second exposures were acquired every two seconds during the elution. Data were reduced at the beamline using the BioXTAS RAW 1.4.0 software (36). The contribution of the buffer to the X-ray scattering curve was determined by averaging frames from the SEC eluent which contained baseline levels of integrated X-ray scattering, UV absorbance, and conductance. Frames were selected as close to the protein elution as possible. If the signal returned to the baseline during the measurement time after protein elution, frames pre- and post-elution were averaged. Otherwise, the baseline was collected only from frames pre-elution. Final q versus I(q) data sets were generated by subtracting the average buffer trace from all elution frames and averaging curves from elution volumes close to the maximum integrated scattering intensity; these frames were statistically similar at both small and large angles. Buffer subtraction, subsequent Guinier fits, and Kratky transformations were done using custom MATLAB (Mathworks) scripts.

### Analytical Ultracentrifugation Sedimentation Velocity (AUC-SV)

Sedimentation velocity experiments were conducted in a ProteomeLab XL-I analytical ultracentrifuge (Beckman Coulter, Indianapolis, IN) following standard protocols unless mentioned otherwise (37,38). Samples in buffer containing 50 mM HEPES pH 7.5, 5 mM DTT and 100, or 200 or 300 mM NaCl were loaded into cell assemblies comprised of double sector charcoal-filled centerpieces with a 12 mm path length and sapphire windows. Buffer density and viscosity were determined in a DMA 5000 M density meter and an AMVn automated micro-viscometer (both Anton Paar, Graz, Austria), respectively. The partial specific volumes and the molecular mass of the protein was calculated based on their amino acid compositions in SEDFIT (https://sedfitsedphat.nibib.nih.gov/software/default.aspx). The cell assemblies, containing identical sample and reference buffer volumes of 390 μL, were placed in a rotor and temperature equilibrated at rest at 20 °C for 2 hours before it was accelerated from 0 to 50,000 rpm. Rayleigh interference optical data were collected at 1-minute intervals for 10 hours. The velocity data were modeled with diffusion-deconvoluted sedimentation coefficient distributions *c(s)* in SEDFIT (https://sedfitsedphat.nibib.nih.gov/software/default.aspx), using algebraic noise decomposition and with signal-average frictional ratio and meniscus position refined with non-linear regression (39). The *s*-values were corrected for time and finite acceleration of the rotor was accounted for in the evaluation of Lamm equation solutions (40). Maximum entropy regularization was applied at a confidence level of P=0.68.

For the sedimentation velocity data of hnRNPA1* samples in the various buffers with increasing salt concentration two-dimensional size-shape distributions, *c*(*s,f/f_0_*) (with the one dimension the *s*-distribution and the other the *ff_0_*-distribution) was calculated with an equidistant *f/f_0_*-grid of 0.25 steps that varies from 0.5 to 3, a linear *s*-grid from 0.5 to 5 S with 100 s-values, and Tikhonov-Phillips regularization at one standard deviation. The velocity data were transformed to *c(s,f/f_0_)*, and *c(s,M)* distributions with *M* the molecular weight, *f/f_0_* the frictional ratio, s the sedimentation coefficient and plotted as contour plots. The color temperature of the contour lines indicates the population of the species (37,38,41).

### Coarse-grained molecular dynamics simulations

#### Initial structure and system preparation

We used Modeller (42) to generate our initial model for the simulations of hnRNPA1* and hSUMO-hnRNPA1* based on the NMR structure of SUMO1 (PDB: 1A5R) (43) and the crystal structure of the RRM1 and RRM2 domains (PDB: lHAl) (44). The LCD* and linker regions were left as an extended coil in the initial structure.

We performed all MD simulations with the MARTINI 3.0.beta.4.17 force-field (http://cgmartini.nl/index.php/martini3beta) (45) using GROMACS 2019.4 (46). The all-atom models of hnRNPA1* and hSUMO-hnRNPA1* were coarse-grained using the Martinize2 python script (47), placed in a cubic box using GROMACS and solvated with the intended NaCl concentration using the Insane python script (48). An elastic network was added to the folded SUMO, RRM1 and RRM2 domains using Martinize2. Interdomain elastic restraints and the elastic network in the LCD* and linker regions were removed. The elastic restraints consisted of a harmonic potential of 500 kJ mol^-1^ nm^-2^ between backbone beads within a 1.2 nm cut-off. Energy minimization was performed for 0.3 ns with a 30 fs timestep using the Berendsen thermostat at 300 K, Berendsen barostat and Verlet cut-off scheme. The system was then equilibrated for 10 ns with a 2 fs timestep using the Velocity-Rescaling thermostat at 300 K, Parinello-Rahman barostat and Verlet cut-off scheme.

#### Molecular dynamics simulations

We first performed coarse-grained MD simulations of hnRNPA1* and hSUMO-hnRNPA1* in which we varied a parameter that modulates the strength of interaction between protein and water. Specifically, we tuned a parameter, *λ*, to rescale the ε-parameter in the Lennard-Jones potential between protein and water beads. We then determined the value of *λ* that gave rise to the best agreement with SAXS data as quantified by the reduced *χ^2^* (*χ^2^r*) between calculated and experimental SAXS profiles. For hnRNPA1*, we chose to tune *λ* at 150 mM NaCl, and performed simulations with *λ* = 1.00, 1.04, 1.06, 1.07, 1.08, 1.09, 1.10 or 1.12 for 10 μs with a 20 fs timestep using the Velocity-Rescaling thermostat at 300 K, Parinello-Rahman barostat and Verlet cut-off scheme, saving conformations every 1 ns. We used Pepsi-SAXS to calculate SAXS profiles from these simulations as previously described (49); specifically we determined the parameters *r*_0_ and *Δρ* as ensemble averages, and *I(0)* and *B* were fitted as free global parameters. These calculations revealed a broad minimum of agreement between experimental and calculated SAXS data for hnRNPA1* in the range *λ* = 1.06 - 1.10. We also performed a comparable analysis using SAXS data at 100 mM NaCl for hSUMO-hnRNPA1* using *λ* = 1.00, 1.02, 1.04, 1.05, 1.06, 1.07, 1.08 or 1.10, revealing a more distinct minimum at *λ* = 1.07, and we chose this value for our further simulations of both hnRNPA1* and hSUMO-hnRNPA1*. We thus performed all our coarse-grained MD simulations of hnRNPA1* and hSUMO-hnRNPA1* with this *λ* = 1.07 and varied the salt concentration to match the experiments (50, 150, 300, 500 or 1000 mM NaCl for hnRNPA1*, and 50, 100, 200, 300, 400 or 500 mM NaCl for hSUMO-hnRNPA1*). We ran these simulations for 20 μs but otherwise with the same parameters as described above.

#### Calculation of SAXS data from simulations

We calculated SAXS data from all-atom models obtained using a modified (49) version of the Backward algorithm (50), in which simulation runs are excluded and energy minimization steps are shortened to 200 steps.

SAXS profiles were calculated from all-atom back-mapped MD trajectories using Pepsi-SAXS 2.4 (51), with experimental SAXS profiles for optimization. Parameters fitted with Pepsi-SAXS are *I(0)*: the forward scattering, *B*: the constant background, *r_0_*: the excluded water volume and *Δρ*: the density of the surface solvent layer.

#### BME reweighting

BME reweighting (52) was performed for simulations with varying ionic strength to refine the ensembles against SAXS data. Fitting of Pepsi-SAXS parameters and BME reweighting was performed as in Larsen et al. 2019 (49), which includes an initial round of BME reweighting to determine optimal weighted ensemble average Pepsi-SAXS parameters, followed by BME reweighting to determine the optimal set of weights for agreement with experimental SAXS. The global scaling parameter *θ* used in the final round of BME reweighting was chosen to obtain a low *χ^2^r* to the experimental SAXS data while retaining a high fraction of effective frames *φ_eff_*. Data points at *q* > 0.2 Å^-1^ were excluded from experimental SAXS data for BME reweighting of the hSUMO-hnRNPA1* ensemble. Our MD simulations, SAXS data and the weights obtained from the reweighting procedure are available at https://github.com/KULL-Centre/papers/tree/master/2020/hnRNPA1-martin-et-al.

#### *R_g_* and contacts

*R_g_* from coordinates was calculated using GROMACS *gyrate* tool. Interdomain contacts between the LCD*, RRMs and hSUMO were calculated using GROMACS *mindist* tool with a 5 Å cut-off and the −*group* flag. For contacts calculations, domains were grouped as residues: (−118)–(−2) (hSUMO), 14-184 (RRMs) and 195–320 (LCD*).

## Results

### Phase separation of hnRNPA1 is highly salt-sensitive

To explore the physicochemical nature of the interactions that mediate LLPS of hnRNPA1, we generated a set of deletion constructs, including the full-length protein (hnRNPA1) and the intrinsically disordered LCD (Fig. 1A). Both constructs were missing a hexapeptide (residues 259-264) that was previously shown to act as a steric zipper and to result in fibrillization from within the dense phase (27). The resulting constructs (denoted with an asterisks) did not undergo fibrillization and were amenable to equilibrium biophysical characterization; this is particularly important when quantifying phase behavior. Full-length hnRNPA1* and the LCD* readily underwent phase separation (Fig. 1B). We have previously reported that the RRMs alone do not phase separate (26).

**Figure 1:**
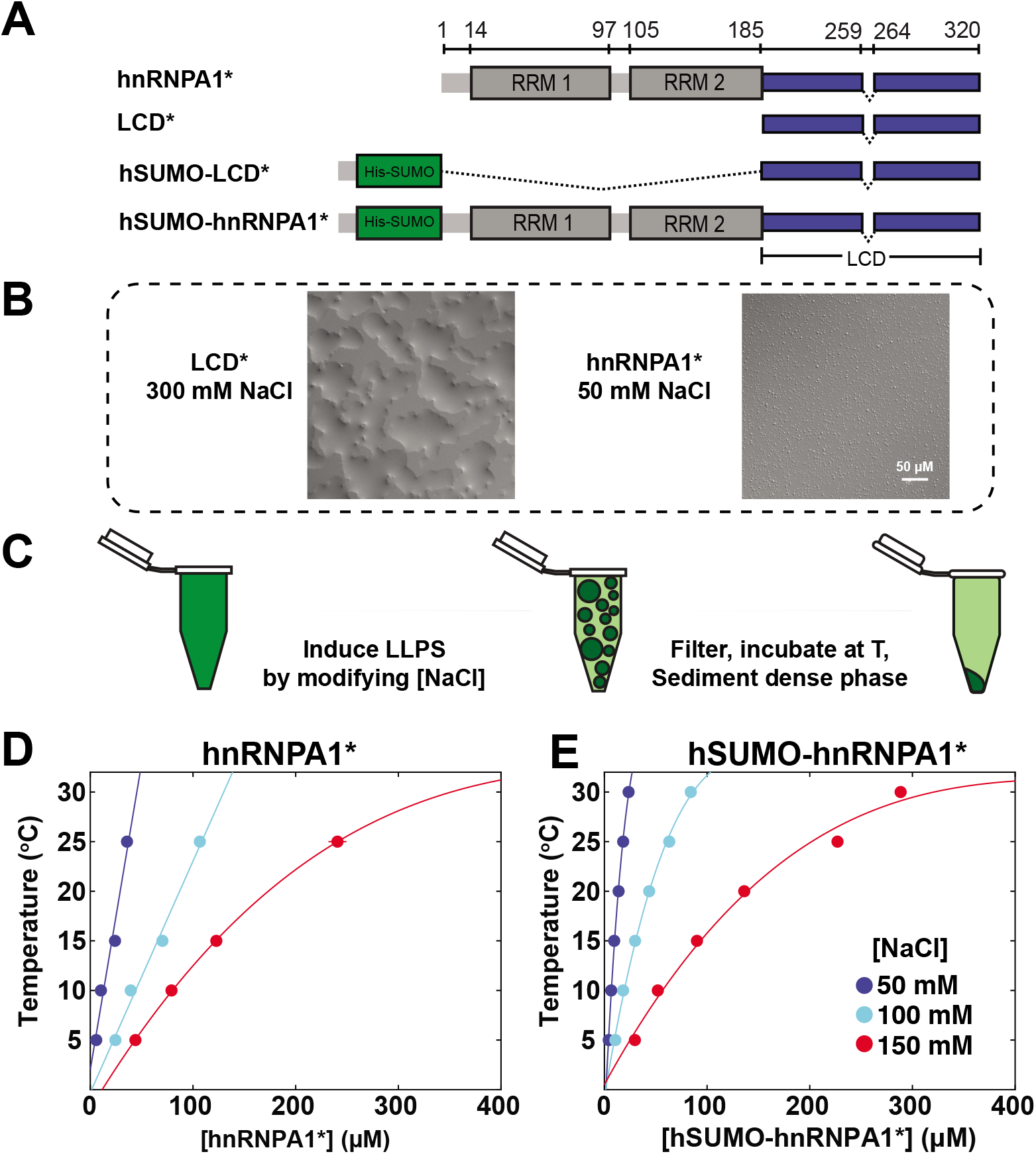
hnRNPA1* undergoes salt-sensitive liquid-liquid phase separation. **(A)** Cartoon schematics for the hnRNPA1* constructs used in this study. hnRNPA1 has two folded RNA binding domains (RRMs 1 and 2, grey), and a C-terminal disordered domain, the so-called low complexity domain (LCD, blue). hSUMO was used for expression and purification and to evaluate interactions of the LCD* with folded domains. A fibrillization-enhancing steric zipper motif (residues 259-264) (27) is removed, indicated by asterisks in the construct name. **(B)** DIC images of hnRNPA1* at 50 mM NaCl and LCD* at 300 mM NaCl. Both proteins are at 300 μM protein concentration. **(C)** Cartoon schematic illustrating the procedure used for mapping the phase diagrams. A stock solution of hnRNPA1 in the one-phase regime is used and LLPS is induced by lowering the NaCl concentration. The phase separated sample is filtered and incubated at the desired temperature. The dense phase is sedimented in a temperature-equilibrated centrifuge, and the light phase concentration determined by UV/Vis spectrophotometric measurement of the light phase (30). **(D)** Phase diagram of full-length hnRNPA1* and **(E)** hSUMO-hnRNPA1* as a function of temperature and NaCl concentration. As NaCl concentration is increased (cool-to-warm color transition), the driving force for phase separation is decreased. The area underneath each curve represents the two-phase regime.

To explore the interactions that mediate phase separation of hnRNPA1 further, we next determined the salt and temperature dependence of the equilibrium dilute phase concentration, *c*_sat_, of full-length hnRNPA1*. From a condition in a high salt storage buffer, in which the protein is highly soluble, phase separation was induced by lowering the NaCl concentration via dilution, and the resulting dense phase was sedimented by centrifugation, resulting in a clear dense phase at the bottom of the tube. The concentration of the dilute phase was determined by UV absorbance measurements (Fig. 1C and Methods) as a function of temperature and NaCl concentration (Fig. 1D) (30). The resulting coexistence curves at different NaCl concentrations are clearly distinct; *c*_sat_ increases with the solution NaCl concentration, demonstrating that phase separation of full-length hnRNPA1* is highly salt-sensitive. The driving force for phase separation increases as salt concentration and temperature decrease. This was also the case for samples of hnRNPA1* tagged with the solubility tag His-SUMO (hSUMO). The temperature dependence implies enthalpically favorable interactions that mediate phase separation, and the salt sensitivity is often interpreted as implicating favorable electrostatic interactions as mediating phase separation.

### Folded domains modulate the solubility of the LCD of hnRNPA1

Given that the LCD of hnRNPA1 is sufficient for phase separation, and phase separation is strongly salt-sensitive as described above, we asked whether the salt-sensitivity is encoded in the LCD of hnRNPA1. The LCD* only contains 10 positively charged and three negatively charged residues in 135 residues, i.e., a fraction of charged residues (FCR) of only 10% and with a predicted net positive charge (Fig. 2A, grey background). This composition is difficult to reconcile with the observation of enthalpically favorable electrostatic interactions mediating phase separation.

**Figure 2:**
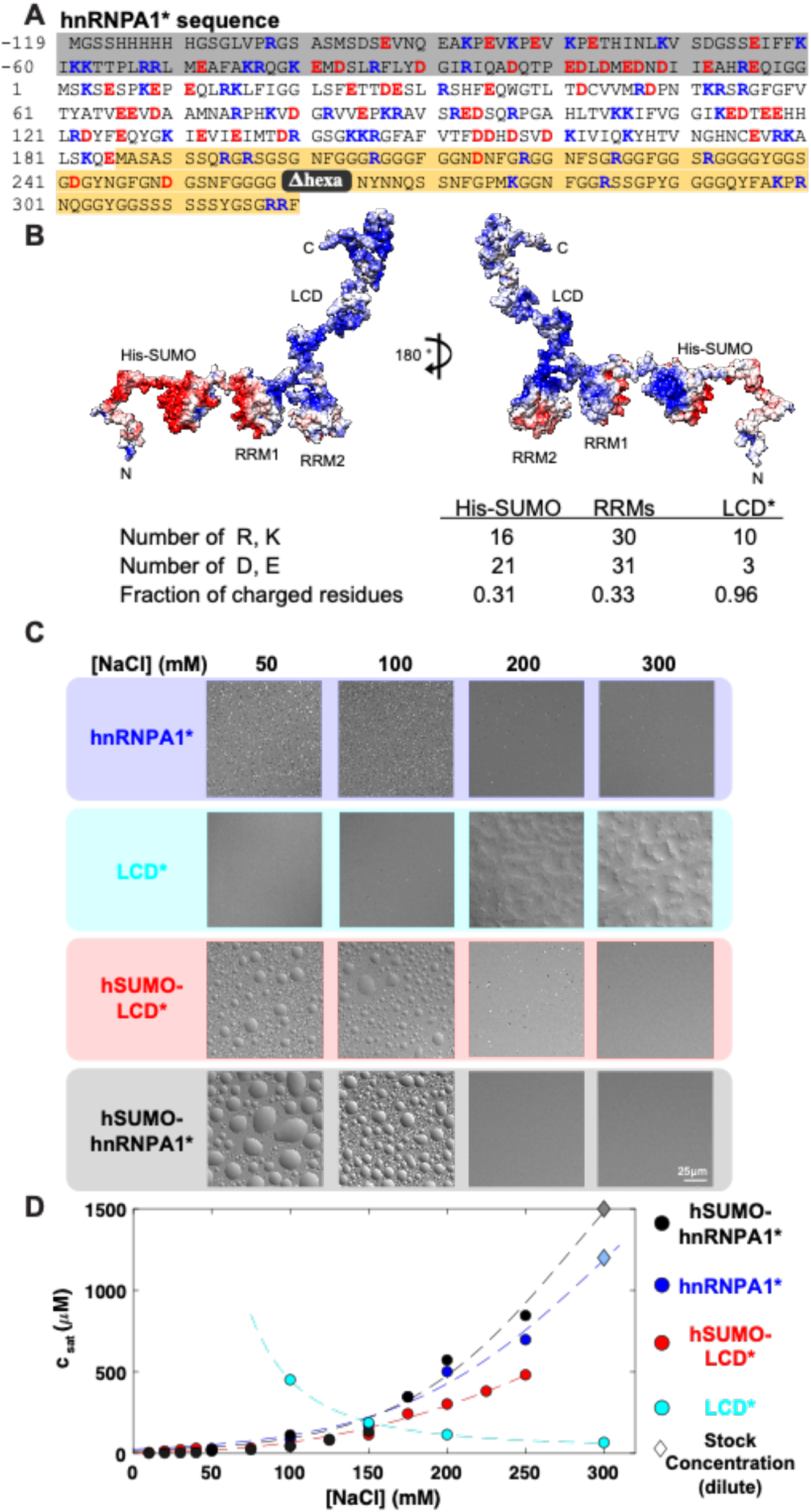
The RRMs modify the phase behavior of hnRNPA1*. **(A)** The sequence of hSUMO-hnRNPA1*; hSUMO residues are shaded in gray. residues 1-185 represent the folded RRM domains, and residues 186-320 represent the disordered LCD (background shaded yellow) and the hexa-peptide that is deleted (259-264) is colored in gray. Positively charged residues are colored blue, negatively charged residues are colored red. The hSUMO tag contains 16 positively charged and 21 negatively charged residues (31% charged). **(B)** Electrostatic potential surface mapped onto the structure of hSUMO-hnRNPA1*; red and blue depict a net negative (−5) and positive (+5) potential, respectively. Chimera was used to generate the surface and APBS surface coloring was automatically generated with default settings. The structure is frame 2574 from MD simulation at 400 mM NaCl. **(C)** DIC images of hnRNPA1*, LCD*, hSUMO-LCD* and hSUMO-hnRNPA1* at 300 μM protein concentration in 50 mM HEPES pH 7.5, 50-300 mM NaCl, 5 mM DTT. Camera settings were identical across each row. Scale bar is 25 μm. The LCD* dense phase wets the glass slide strongly. **(D)** Dilute phase concentrations of full-length hnRNPA1 (black) and the LCD (red) as a function of NaCl concentration at 20°C. Diamonds represent the stock concentrations of samples that did not phase separate at 20°C. The dashed lines are a fit to a logistic function to guide the eye.

This raised the question whether there may be salt-sensitive interactions in or between other domains of hnRNPA1*. The folded RRMs (Fig. 2A, white background) have a higher fraction of charged residues than the LCD*. Displaying the charge distribution on the surface of the structure of the hnRNPA1 RRMs (Fig. 2B) revealed two distinct faces; one face is predominantly negatively charged (Fig. 2B, red) and the other face is predominantly positively charged (blue); the latter binds the negatively charged RNA. (hnRNPA1* was expressed with a proteolytically cleavable hSUMO solubility tag, which also has a higher fraction of charged residues and a negatively charged surface (Fig. 2B and below).) These observations lead us to hypothesize that the salt-sensitivity of hnRNPA1* phase behavior may be the result of its folded domains or interactions between them and the LCD*.

We thus determined the salt dependence of *c*sat of the LCD*, which we found to be opposite to that of full-length hnRNPA1* (Fig. 2C,D). *c*sat decreased with increasing NaCl concentration, indicating a stronger driving force for phase separation at higher salt concentrations. These observations are in agreement with the expectation from our previous work that phase separation of the LCD is driven by interactions among aromatic residues uniformly distributed along the sequence (17). The LCD of hnRNPA2 has also been reported to phase separate more strongly with increasing salt concentration (53). Hence, the addition of the folded RRMs reversed the salt dependence of LCD phase separation.

To test whether this effect was specific to the folded RRMs, we took advantage of the hSUMO affinity/solubility tag that we used to express and purify hnRNPA1* constructs (Fig. 2A). SUMO is a small ubiquitin-like modifier with positive and negative surfaces similar to the RRMs and net negative charge (Fig. 2B). The salt-dependence of hSUMO-LCD* phase behavior was nearly identical to that of full-length hnRNPA1*, suggesting that the effect of either folded domain module, the RRMs or hSUMO, on the LCD* may stem from similar interactions or from the modulation of the LCD* solubility by the folded domains. Adding both folded domain modules to generate hSUMO-hnRNPA1* (i.e., including hSUMO and the RRMs) increased the solubility even more at the higher salt concentrations. In fact, hSUMO-hnRNPA1* lost the ability to undergo phase separation past 250 mM NaCl (Fig. 2C, D; tested at protein concentrations of up to ~1200 μM), suggesting an additive effect of the folded domains on solubility.

Interestingly, at NaCl concentrations up to 175 mM NaCl, hSUMO-hnRNPA1* had a slightly stronger driving force for phase separation than hnRNPA1* or hSUMO-LCD* as indicated by the lower values of *c*sat. These results suggest that electrostatic interactions between the folded domains and the LCD* contribute to phase separation at NaCl concentrations that do not screen these.

Taken together, these data support the hypothesis that electrostatic interactions between folded domains and the LCD of hnRNPA1 contribute to phase separation at low ionic strengths, that they are disrupted at high ionic strength, and that folded domains solubilize the LCD at these higher ionic strengths. Folded domains with charged surfaces are indeed expected to have increased solubility with increasing NaCl concentrations until salting-out is favored at high ionic strength (54).

### The global dimensions of hnRNPA1* report on salt-sensitive intramolecular interactions

We reasoned that the transition of hnRNPA1* from salt concentrations that enable enhanced phase separation to those that solubilize hnRNPA1* should be readily observable by measurements of single-chain dimensions. We expected that the increased solubility at increasing NaCl concentrations would be accompanied by expanding protein conformations. We carried out size exclusion chromatography-coupled SAXS (SEC-SAXS) measurements at a variety of NaCl concentrations (Fig. S1-3). The SAXS data of fulllength hnRNPA1* showed little change between 50 and 400 mM NaCl (Fig. 3), but hSUMO-hnRNPA1* showed a clear increase in its global dimensions (Fig. 4).

**Figure 3:**
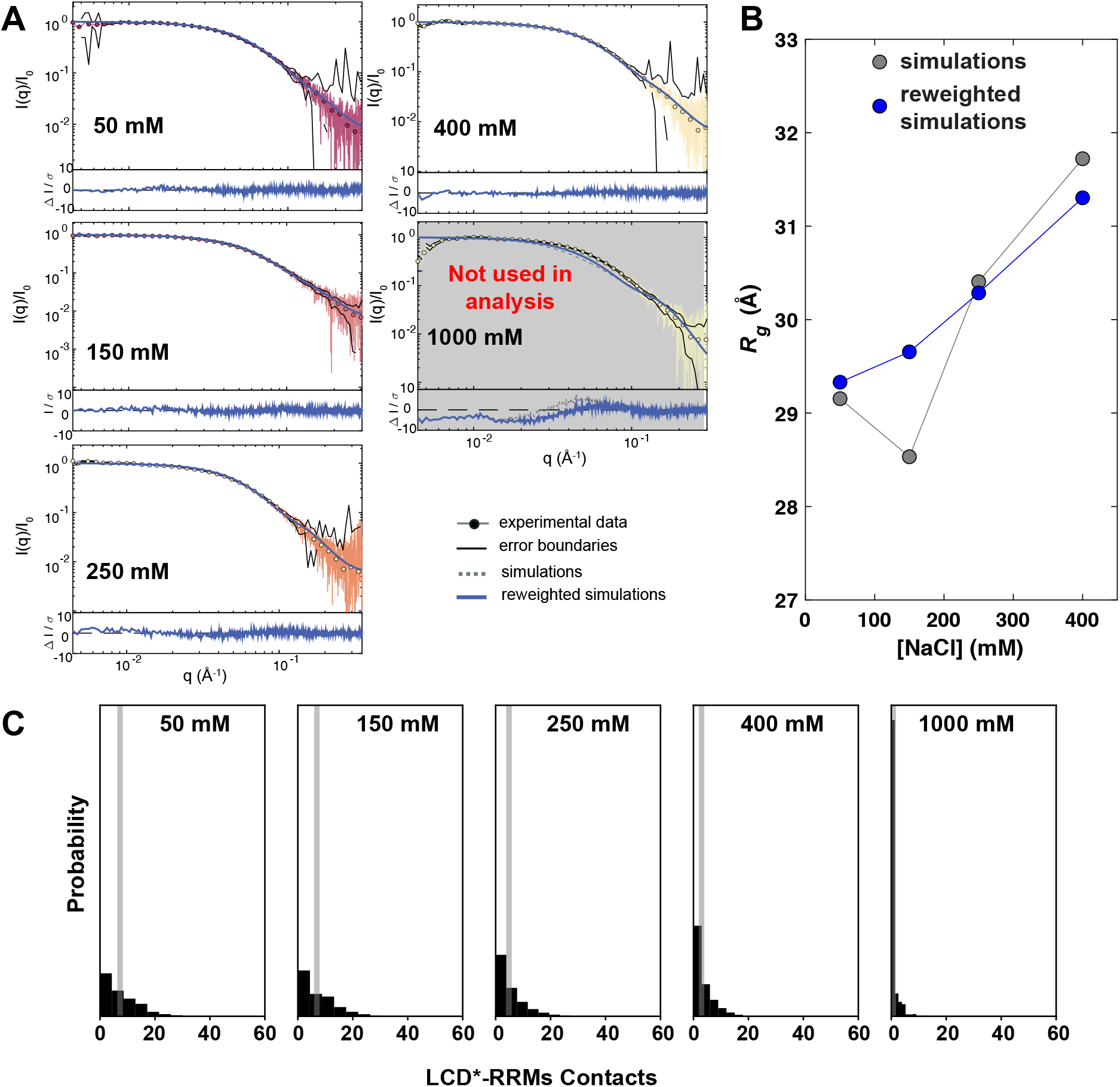
hnRNPA1* shifts equilibrium between compact and extended conformations in a salt concentration-dependent manner. **(A)** SEC-SAXS data acquired of hnRNPA1 in 50 mM Tris pH 7.5, 10 mM DTT, 2 mM TCEP and 50 mM NaCl - 1000 mM NaCl. The fit of calculated data from simulations, before and after reweighting, to the experimental data is shown. **(B)** *R*_g_ values calculated from simulations as a function of NaCl concentration. **(C)** Increasing NaCl concentrations progressively reduce the number of RRM-LCD interactions; average number shown as grey bar.

**Figure 4:**
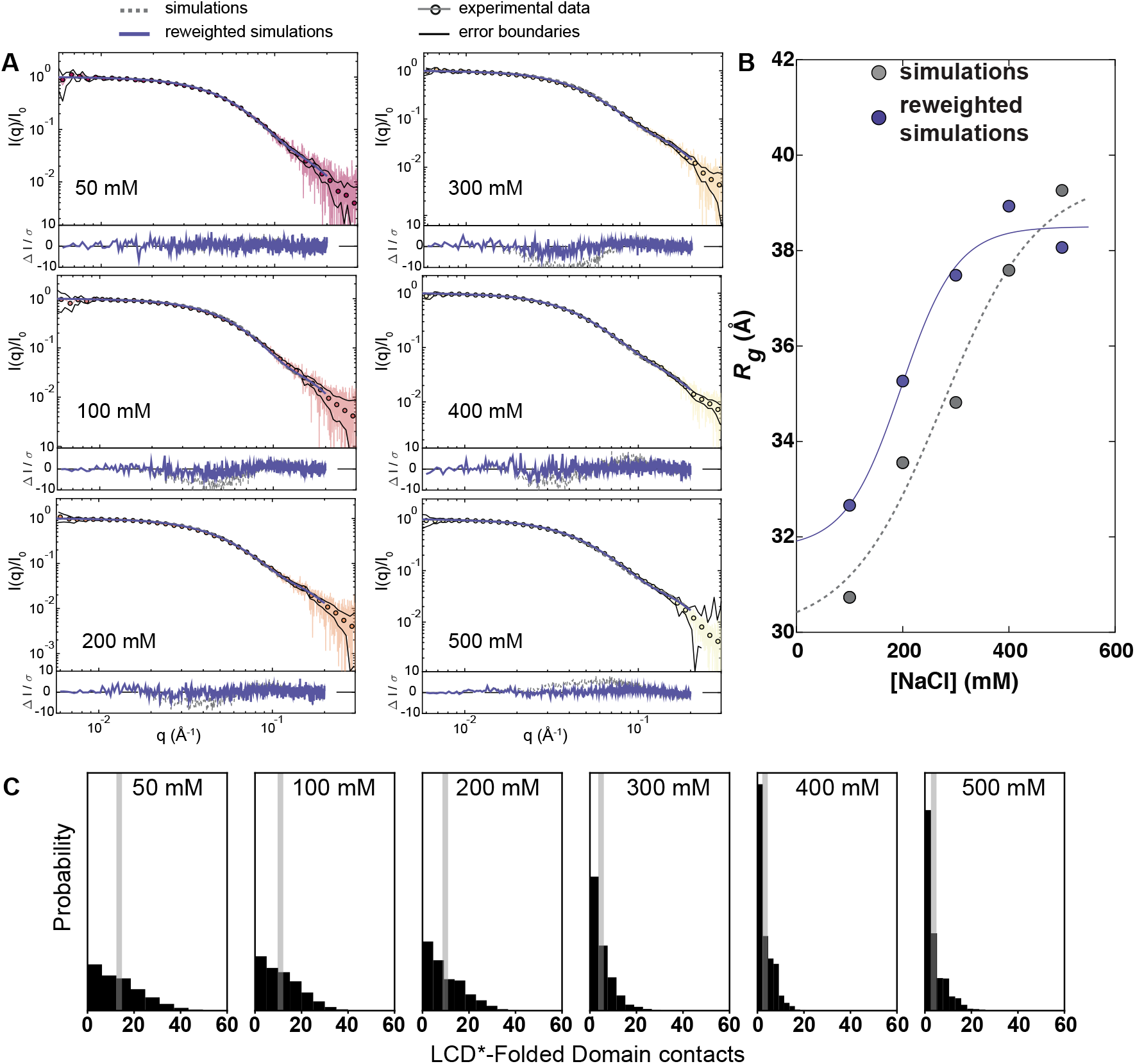
hSUMO-hnRNPA1* shifts equilibrium between compact and extended conformations in a salt concentration-dependent manner. **(A)** SEC-SAXS data of hSUMO-hnRNPA1* in 50 mM Tris pH 7.5, 10 mM DTT, 2 mM TCEP and 50 mM NaCl - 500 mM NaCl. The fit of calculated data from simulations, before and after reweighting, to the experimental data is shown. Guinier analysis plots are shown in insets. **(B)** *R*_g_ values calculated from simulations as a function of NaCl concentration. The data was fit to logistic functions that indicate minimum dimensions at low NaCl concentration, a maximum extension at high NaCl concentration and a transition with a midpoint at ~175 mM NaCl, suggesting screening of interactions between oppositely charged regions of hSUMO-hnRNPA1*. Simulations recapitulate the trend and after reweighting show close agreement. **(C)** Increasing NaCl concentrations progressively reduce the number of RRM-LCD* and hSUMO-LCD* interactions; average number shown as grey bar.

We turned to coarse-grained MD simulations with the MARTINI forcefield to obtain a molecular picture of the potential interdomain interactions that affect the compaction of hnRNPA1* and hSUMO-hnRNPA1*. Focusing first on hnRNPA1*, we initially performed simulations at 150 mM NaCl. Comparing the resulting conformational ensemble with the SAXS data, however, revealed it was somewhat too compact (Fig. S4). We have previously demonstrated that rescaling of protein-water interactions may be used to alleviate this issue and results in conformational ensembles of RNA-binding proteins that agree well with experiment (49). We thus changed the protein-water interaction by tuning a parameter, λ, that increases the interaction between protein and water, and chose the value that resulted in the best fit between experimental and calculated SAXS data for both hnRNPA1* and hSUMO-hnRNPA1* (Fig. S4 and Methods). We find that an increase of 7% (λ=1.07) is sufficient to obtain a good fit to the data, and note that we previously obtained a very similar value (λ=1.06) on a different protein (49), suggesting that the magnitude might be general and transferable. We thus proceeded to use λ=1.07 for coarse-grained simulations at the NaCl concentrations used for SAXS measurements.

The simulations reveal a small salt-dependent expansion of hnRNPA1* and were generally in good agreement with the SAXS data between 50 – 400 mM NaCl and resulted in low values of χ^2^ (Fig. S4C), but the calculated SAXS curves showed small, non-random deviations from experiment (Fig. 3A). The simulations at 1000 mM NaCl agreed less well with the experiments, suggesting that the MARTINI model used does not capture the effect of such high salt concentrations well. We have previously demonstrated that reweighting against experiments is robust as long as the initial simulation is relatively good (49). We thus used a Bayesian/maximum entropy approach (52,55) to reweight the conformational ensembles of hnRNPA1* against the experimental SAXS data (Fig. 3A,B, Fig. S4,5), and to obtain an improved fit to the experimental SAXS data at all NaCl concentrations, thus generating ensembles that are in full accordance with the experiments. Because reweighting is most robust when the simulations are in reasonably good agreement before reweighting, we focused our analysis on the range between 50 – 400 mM NaCl. The ensemble *R*_g_ is ~29 Å at 50 mM NaCl, increases slightly with increasing NaCl concentration and reaches 31 Å at 400 mM NaCl. These observations are in agreement with a scenario in which weak, transient intramolecular electrostatic interactions compact hnRNPA1*; these interactions are screened at increasing ionic strength, which results in an expansion of hnRNPA1* and a maximal average expansion at NaCl concentrations above the physiological range. Notably, the salt sensitivity of the *Rg* occurs in the same concentration window as the strongest change of the saturation concentration.

Inspecting the resulting conformational ensembles showed extensive RRM-LCD* interactions at low NaCl concentrations which were progressively disrupted at increasing concentrations (Fig. 3C). The simulations thus suggest that increasing salt concentrations shift the equilibrium between conformations in which the LCD* associates with the RRMs and conformations in which the LCD* is liberated from the RRMs. We note that a similar picture was obtained analyzing the ensembles prior to reweighting, demonstrating that the MARTINI force field itself captured the observed salt dependency of the interactions.

We used the same simulation approach for hSUMO-hnRNPA1* and found a bigger increase of its dimensions with increasing NaCl concentration in agreement with the experimental SAXS data (Fig. 4A). Again, we used reweighting against the SAXS data to improve agreement further (Fig. 4A,B, Fig. S4) and analyzed the resulting ensembles for interactions between the LCD* and both the RRMs and hSUMO. As for hnRNPA1*, we find extensive and salt-dependent interactions between LCD* and the RRMs, as well as interactions between LCD* and hSUMO (Fig. 4C).

As an independent means of validating the expansion of hSUMO-hnRNPA1* we performed two-dimensional analytical ultracentrifugation sedimentation velocity (AUC-SV) analysis. The results show that the shape factor, *f/f_0_*, also becomes larger with increasing NaCl concentration, which signifies an increase in extended shape and supports the idea that the LCD* is released from its interactions with the folded domains and increases the hydrodynamic drag of hSUMO-hnRNPA1* (Fig. S6).

The SAXS data, AUC-SV analysis and simulation data all support the existence of electrostatic interactions of the LCD*s with the folded domains of hnRNPA1* and hSUMO-hnRNPA1*. To examine whether these changes are somehow driven by a salt-dependent expansion of the LCD*, we also determined SAXS profiles for LCD* samples (Fig. 5). Normalized Kratky plots reveal a slight compaction of the LCD* with increasing NaCl (Fig. 5A), in agreement with stronger aromatic-aromatic interactions, and with the fact that the solubility of the LCD* alone decreased with increasing NaCl concentration. We analyzed these SAXS profiles using an empirical molecular form factor, indeed showing a decrease of *R*_g_ and the scaling exponent *ν*_app_ (Fig. 5B). Thus, it appears that the salt-dependent change in dimensions of hnRNPA1* and in particular hSUMO-hnRNPA1* are not driven by an intrinsic expansion of the LCD*, but rather happens over a concentration range where the LCD* compacts. These results point to a partial cancelation of the expected expansion of hnRNPA1* and hSUMO-hnRNPA1* due to the release of RRM-LCD* and hSUMO-LCD* interactions, and suggest that the observed change in overall compaction of these two constructs might underestimate the effect of the release of interactions with the LCD*.

**Figure 5:**
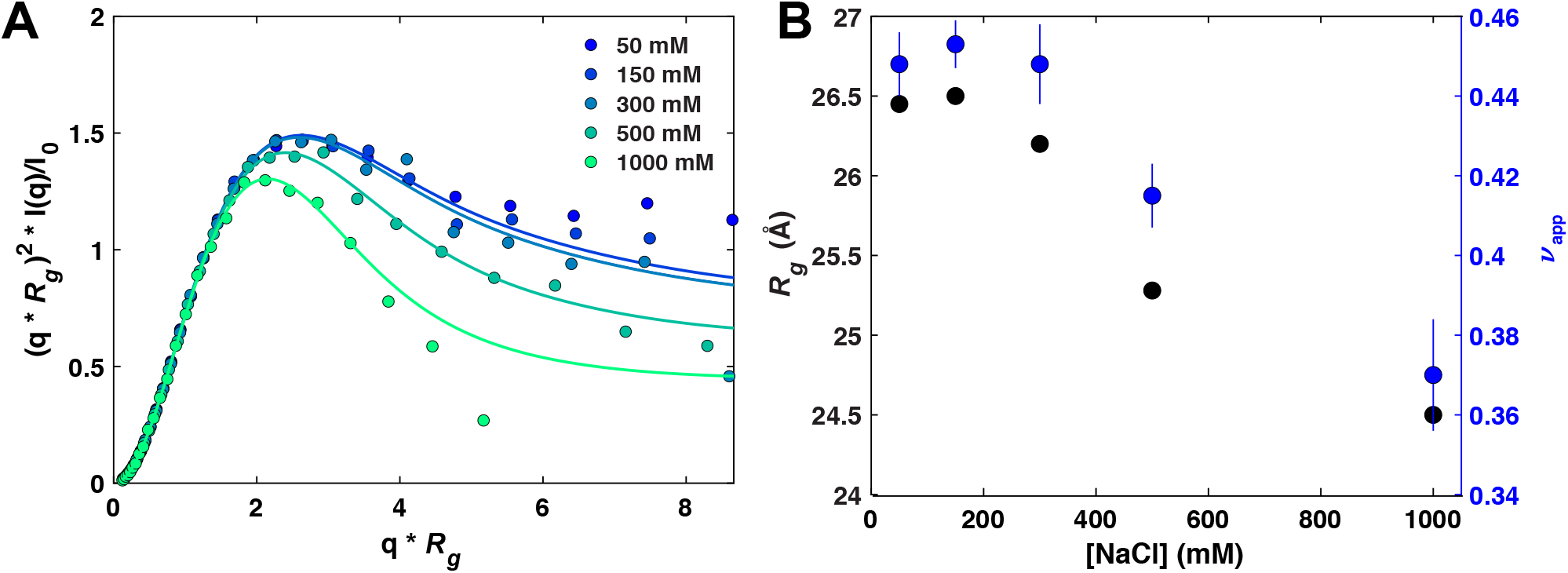
The LCD* compacts with increasing salt concentration. **(A)** Normalized Kratky plots of LCD* samples in different NaCl concentrations. Data was recorded by SEC-SAXS with a co-flow cell. **(B)** Radii of gyration and scaling exponents (v_app_, as a measure of size) as a function of NaCl concentration as analyzed using a molecular form factor.

Having established transient, salt-dependent interactions between the LCD* and the different folded domains, we used the reweighted simulations to analyze which residues form these interactions. We thus calculated the number of contacts formed between the LCD* and the RRMs in both hnRNPA1* and hSUMO-hnRNPA1*, as well as the LCD*-hSUMO interactions in hSUMO-hnRNPA1* (Fig. 6). Comparing the ensembles of hnRNPA1* and hSUMO-hnRNPA1* we find that the same residues in both the LCD* and RRMs make interactions in both constructs (Fig. 6A,B). It also appears that at 50 mM NaCl the interactions between the LCD* and the RRMs are strengthened by the presence of and interactions with hSUMO. We find many of the contacts from the LCD* to the RRMs on the residues opposite of the RNA-binding face of hnRNPA1* but also extending to its edge (Fig. 6D). This finding suggests that the net positively charged LCD* interacts transiently with the negatively charged face of the folded RRMs at low NaCl concentrations. However, some interactions are also made with the positively charged face of the RRMs, highlighting that the LCD-RRM interactions are not purely electrostatic in nature even if these dominate.

**Figure 6:**
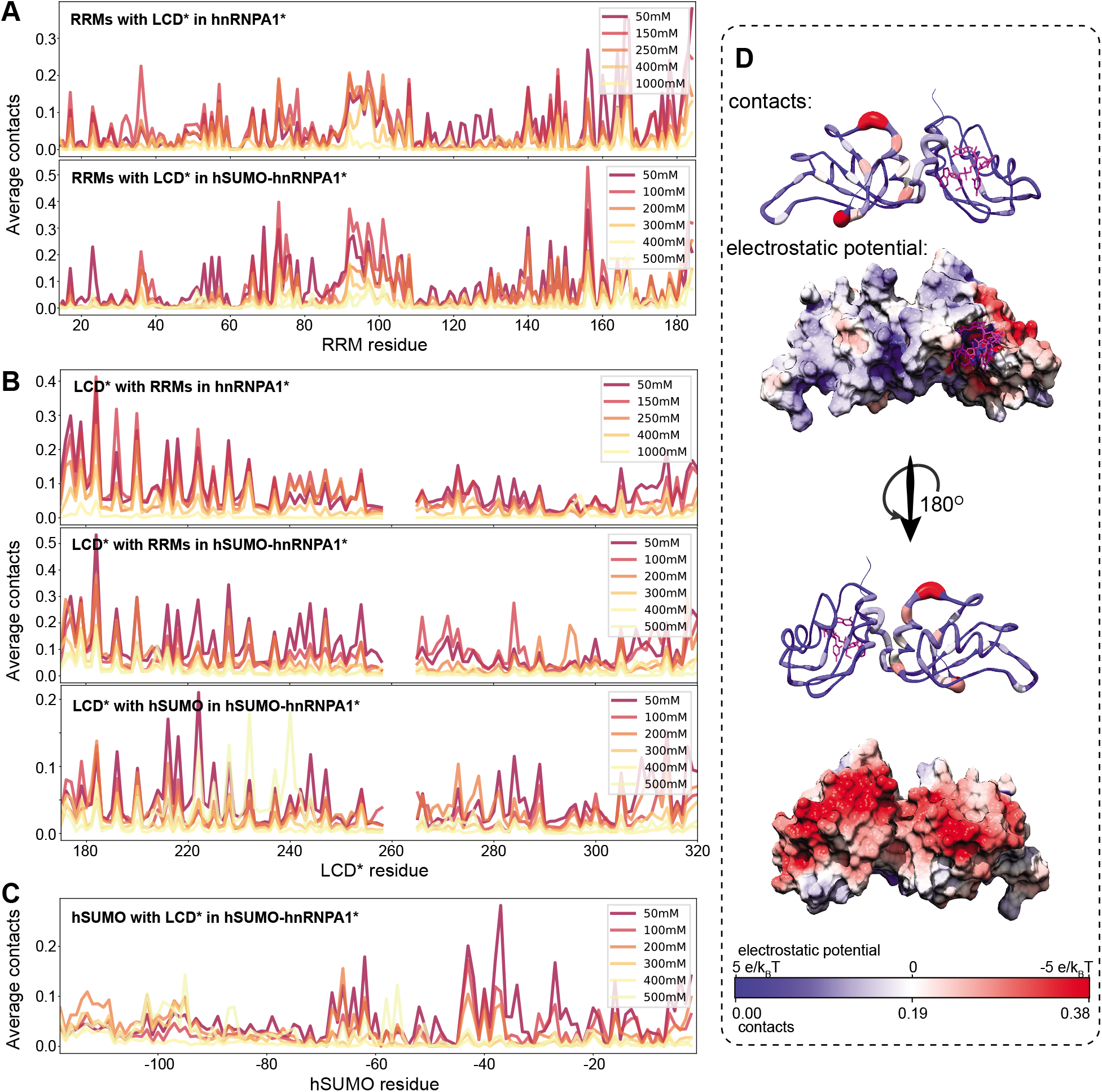
The LCD* interacts with the folded domains in a salt-sensitive manner. **(A)** Average contact number of the LCD* on the sequence of the RRMs. **(B)** Average contact number of the folded domains on the sequence of the LCD*. **(C)** Average contact number of the LCD* on the sequence of hSUMO. The y-axis scales differ between plots. **(D)** Contacts from LCD* on the RRMs displayed in sausage representation with the number of contacts colored from blue to red and the electrostatic potential surface of the RRMs colored from red to blue indicating negative to positive. Bound RNA is shown in purple in the sausage plot. PDB: 4YOE (28).

## Discussion

Disordered LCDs of RNA-binding proteins are often alone sufficient for mediating phase separation, but how they function in the context of the full protein and how the folded domains modulate LCD phase behavior has remained incompletely understood. Here, we show that phase separation of hnRNPA1 is salt-sensitive while phase separation of the LCD alone is promoted with increasing ionic strength. We find that folded domains play dual roles in modulating LCD phase behavior: (1) Electrostatic interactions between folded domains and the disordered LCD can contribute to phase separation at low ionic strength; increasing ionic strengths screen these interactions; (2) at higher ionic strengths, the folded domains solubilize the fusion protein (Fig. 7).

**Figure 7:**
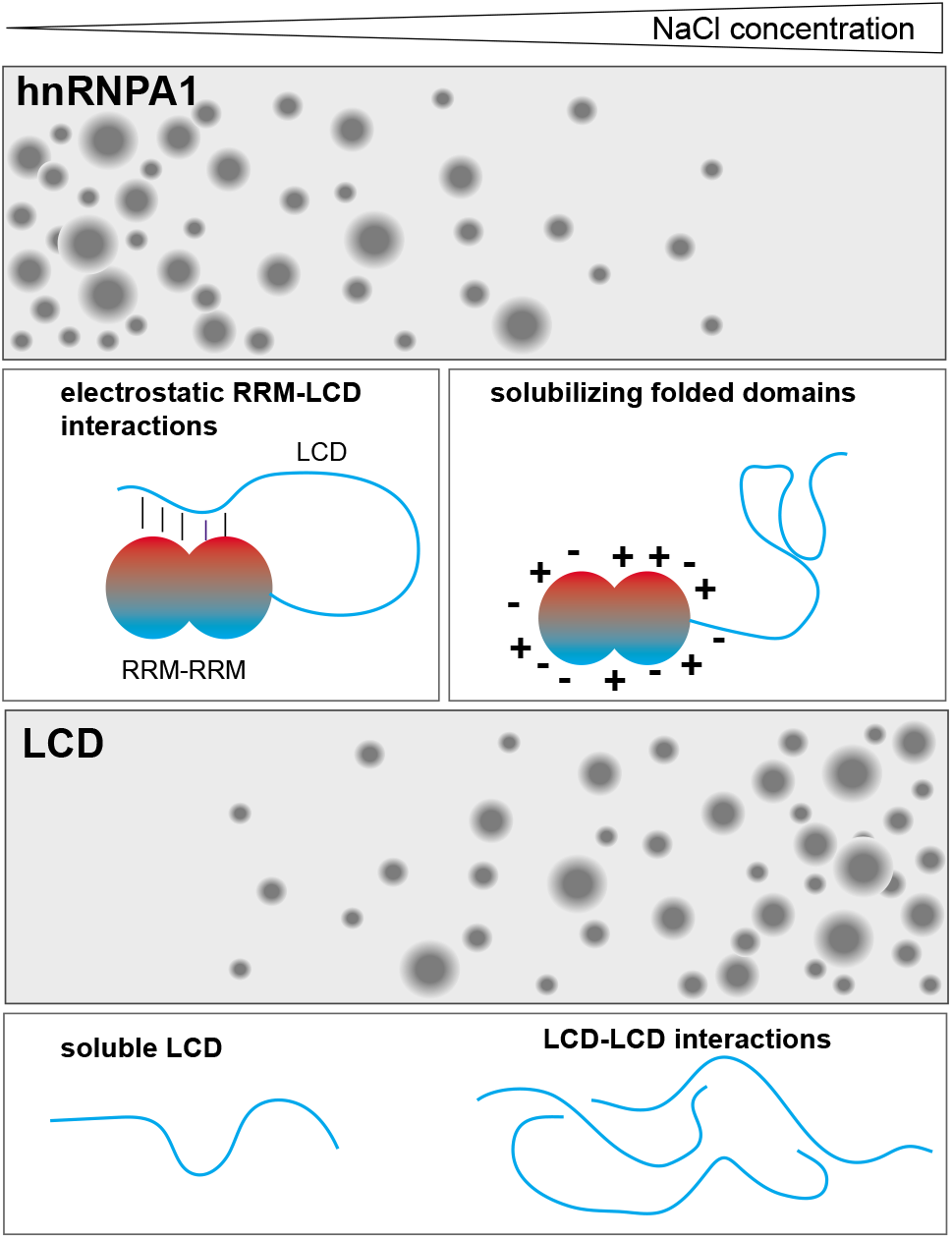
Interactions between the folded RRMs and the disordered LCD modulate the phase behavior of hnRNPA1. The LCD undergoes LLPS even at high salt concentrations. The electrostatic interactions between the RRMs and the LCD contribute to LLPS of full-length hnRNPA1. They are screened with increasing salt concentration and the resulting expanded conformation is more soluble than the LCD alone. This mechanism highlights the ability of the LCD to modulate phase behavior of hnRNPA1, e.g. also via post-translational modifications.

Typical LCDs of the type found in hnRNPA1 and related proteins are polar tracts interspersed with aromatic and few charged residues (15). The main driving force for their phase separation seems to stem from a combination of aromatic-aromatic and aromatic-arginine interactions (13,15,17,20,56), which are compatible with high ionic strength. This is in agreement with our observation that phase separation of the hnRNPA1 LCD is promoted at high salt concentration. In contrast, we did not observe phase separation of full-length hnRNPA1 at NaCl concentrations above 200 mM, even at millimolar protein concentrations. This observation suggested the presence of salt-sensitive electrostatic interactions between the LCD and the RRMs that modulate phase behavior. Indeed, we observed intramolecular interactions between the LCD and the RRMs by coarse-grained MD simulations, and between the LCD and hSUMO in a fusion protein construct. We further showed that the global dimensions of full-length hSUMO-hnRNPA1 expand concomitantly with increasing salt concentration, caused by screening electrostatic interactions that stabilize compact conformations involving LCD-RRM and LCD-hSUMO contacts.

The enhanced driving force for phase separation of full-length hnRNPA1 as compared to the LCD at low salt concentration is in agreement with a model in which the RRM-LCD interactions can occur in *trans* and contribute to phase separation (Fig. 6). As the salt concentration increases and the RRM-LCD interactions are screened, the saturation concentration increases massively, to the point that we were unable to detect phase separation. Our and other previously published data point to the solubilizing influence of the folded domains. Under conditions where the LCD does not associate with the RRMs, the excluded volume of the RRMs decreases the potential for LCD-LCD mediated phase separation and the higher charge content of the RRMs has a solubilizing effect. hSUMO has a similar effect and adding both folded domain modules compounds the effect. In fact, this solubilizing property of folded domains has been used to prevent phase separation of strongly self-associating LCDs until the solubilizing domain is proteolytically cleaved (21,22,57,58). The solubilizing effect of karyopherins on FUS and other RNA-binding proteins (23,59–61) may work through similar mechanisms. We suggest that solubilization is the result of salting in, which derives from a combination of several molecular effects, including ion accumulation around the folded domains (54,62).

A key result is that solubilization through folded domains and their potential interactions with LCDs may obscure the physicochemical nature of the interactions that mediate LCD phase separation. We concluded from domain deletion experiments that the LCD was necessary and sufficient for phase separation. The salt sensitivity of hnRNPA1 phase behavior could have further led us to conclude that electrostatic interactions were key drivers of phase separation (and in fact did in (26)). However, a careful analysis of the phase behavior of the LCD alone and of different domain constructs confirmed that the interactions underlying LCD phase separation are hydrophobic in nature, that electrostatic RRM-LCD interactions contribute to phase behavior at low salt, and that the RRMs are solubilizing at high salt. These insights demand a careful assessment of the physicochemical interactions underlying phase behavior of multi-domain proteins. They should also be taken into account when phase-separating proteins are fused to reporter proteins.

Our experiments were carried out in the absence of RNA and therefore lead us to a conceptual understanding of the influence of folded domains on LCD phase behavior in vitro, not necessarily of hnRNPA1 biology in cells. However, it is highly relevant for considerations of the effect of fusion proteins in cells. Also, the interplay between the folded RNA-binding domains and the disordered LCD suggests an interesting mechanism for tuning hnRNPA1 phase separation in cells. RNA binding and posttranslational modifications are likely to modulate the attractive RRM-LCD interactions, and several independent inputs may thus be able to modulate the saturation concentration. RNA may screen the positive face of the RRMs and create more negative surfaces for interactions with the LCD. It has recently been demonstrated that G3BP1 encodes interactions between several domains and that their modulation by posttranslational modifications tunes the sensitivity of G3BP1 to the levels of exposed mRNA in cells and therefore adjusts its driving force for stress granule assembly (63,64). In Npm1, the main protein that mediates the formation of the Granular Component of the nucleolus via phase separation, intramolecular interactions that modulate compaction and driving force for phase separation have also been observed (65). TIA1, another RNA-binding protein with roles in the regulation of transcription, splicing and the stress response, also shows interactions between tandem repeats of RRMs and its LCD (66) although the functional consequences of this are unknown. These results and our current report suggest that the threshold concentration for phase separation of RNA-binding proteins may not be a fixed value in cells but can be tuned depending on inputs from signaling pathways.

Modulating interactions between folded domains and LCDs may constitute a general principle for tuning the phase behavior of proteins in response to input signals. Further work is needed to disentangle the multitude of protein-protein and protein-RNA interactions and their balance in mediating phase separation under physiological conditions.

## Supplementary Data

Supplementary Data are available.

## ACKNOWLEDGEMENT

We are grateful to Max Frenkel for technical help and to Jill Bouchard, Melissa Marzahn, Ivan Peran, Richard Kriwacki, Irina Ritsch, Elisabeth Lehmann, Gunnar Jeschke and Frédéric Allain for fruitful discussions. We thank Yong Wang, Francesco Pesce and Giulio Tesei for help with and input to running and analyzing the MD simulations. We thank Victoria Frohlich, Aaron Pitre, Jennifer Peters, and Sharon King for help with microscopy. We acknowledge the use of the Molecular Interaction Analysis facility in Protein Technologies Center in the Department of Structural Biology at SJCRH for analytical ultracentrifugation experiments. We thank Srinivas Chakravarthy and the BioCAT beamline staff at the Advanced Photon Source for assistance with SAXS measurements.

## REFERENCES

1. Shin, Y. and Brangwynne, C.P. (2017) Liquid phase condensation in cell physiology and disease. Science, 357.

2. Banani, S.F., Lee, H.O., Hyman, A.A. and Rosen, M.K. (2017) Biomolecular condensates: organizers of cellular biochemistry. Nat Rev Mol Cell Biol, 18, 285–298.

3. Boeynaems, S., Alberti, S., Fawzi, N.L., Mittag, T., Polymenidou, M., Rousseau, F., Schymkowitz, J., Shorter, J., Wolozin, B., Van Den Bosch, L. et al. (2018) Protein Phase Separation: A New Phase in Cell Biology. Trends Cell Biol, 28, 420–435.

4. Strom, A.R., Emelyanov, A.V., Mir, M., Fyodorov, D.V., Darzacq, X. and Karpen, G.H. (2017) Phase separation drives heterochromatin domain formation. Nature, 547, 241–245.

5. Larson, A.G., Elnatan, D., Keenen, M.M., Trnka, M.J., Johnston, J.B., Burlingame, A.L., Agard, D.A., Redding, S. and Narlikar, G.J. (2017) Liquid droplet formation by HP1alpha suggests a role for phase separation in heterochromatin. Nature, 547, 236–240.

6. Gibson, B.A., Doolittle, L.K., Schneider, M.W.G., Jensen, L.E., Gamarra, N., Henry, L., Gerlich, D.W., Redding, S. and Rosen, M.K. (2019) Organization of Chromatin by Intrinsic and Regulated Phase Separation. Cell, 179, 470–484 e421.

7. Sabari, B.R., Dall’Agnese, A., Boija, A., Klein, I.A., Coffey, E.L., Shrinivas, K., Abraham, B.J., Hannett, N.M., Zamudio, A.V., Manteiga, J.C. et al. (2018) Coactivator condensation at superenhancers links phase separation and gene control. Science, 361.

8. Boija, A., Klein, I.A., Sabari, B.R., Dall’Agnese, A., Coffey, E.L., Zamudio, A.V., Li, C.H., Shrinivas, K., Manteiga, J.C., Hannett, N.M. et al. (2018) Transcription Factors Activate Genes through the Phase-Separation Capacity of Their Activation Domains. Cell, 175, 1842–1855 e1816.

9. Boehning, M., Dugast-Darzacq, C., Rankovic, M., Hansen, A.S., Yu, T., Marie-Nelly, H., McSwiggen, D.T., Kokic, G., Dailey, G.M., Cramer, P. et al. (2018) RNA polymerase II clustering through carboxy-terminal domain phase separation. Nat Struct Mol Biol, 25, 833–840.

10. Su, X., Ditlev, J.A., Hui, E., Xing, W., Banjade, S., Okrut, J., King, D.S., Taunton, J., Rosen, M.K. and Vale, R.D. (2016) Phase separation of signaling molecules promotes T cell receptor signal transduction. Science, 352, 595–599.

11. Case, L.B., Zhang, X., Ditlev, J.A. and Rosen, M.K. (2019) Stoichiometry controls activity of phase-separated clusters of actin signaling proteins. Science, 363, 1093–1097.

12. Li, P., Banjade, S., Cheng, H.C., Kim, S., Chen, B., Guo, L., Llaguno, M., Hollingsworth, J.V., King, D.S., Banani, S.F. et al. (2012) Phase transitions in the assembly of multivalent signalling proteins. Nature, 483, 336–340.

13. Nott, T.J., Petsalaki, E., Farber, P., Jervis, D., Fussner, E., Plochowietz, A., Craggs, T.D., Bazett-Jones, D.P., Pawson, T., Forman-Kay, J.D. et al. (2015) Phase transition of a disordered nuage protein generates environmentally responsive membraneless organelles. Mol Cell, 57, 936–947.

14. Mittag, T. and Parker, R. (2018) Multiple Modes of Protein-Protein Interactions Promote RNP Granule Assembly. J Mol Biol, 430, 4636–4649.

15. Wang, J., Choi, J.M., Holehouse, A.S., Lee, H.O., Zhang, X., Jahnel, M., Maharana, S., Lemaitre, R., Pozniakovsky, A., Drechsel, D. et al. (2018) A Molecular Grammar Governing the Driving Forces for Phase Separation of Prion-like RNA Binding Proteins. Cell, 174, 688–699 e616.

16. Martin, E.W. and Mittag, T. (2018) Relationship of Sequence and Phase Separation in Protein Low-Complexity Regions. Biochemistry, 57, 2478–2487.

17. Martin, E.W., Holehouse, A.S., Peran, I., Farag, M., Incicco, J.J., Bremer, A., Grace, C.R., Soranno, A., Pappu, R.V. and Mittag, T. (2020) Valence and patterning of aromatic residues determine the phase behavior of prion-like domains. Science, 367, 694–699.

18. Harmon, T.S., Holehouse, A.S., Rosen, M.K. and Pappu, R.V. (2017) Intrinsically disordered linkers determine the interplay between phase separation and gelation in multivalent proteins. Elife, 6.

19. Lin, Y.H., Forman-Kay, J.D. and Chan, H.S. (2016) Sequence-Specific Polyampholyte Phase Separation in Membraneless Organelles. Phys Rev Lett, 117, 178101.

20. Lin, Y., Currie, S.L. and Rosen, M.K. (2017) Intrinsically disordered sequences enable modulation of protein phase separation through distributed tyrosine motifs. J Biol Chem, 292, 19110–19120.

21. Burke, K.A., Janke, A.M., Rhine, C.L. and Fawzi, N.L. (2015) Residue-by-Residue View of In Vitro FUS Granules that Bind the C-Terminal Domain of RNA Polymerase II. Mol Cell, 60, 231–241.

22. Wang, A., Conicella, A.E., Schmidt, H.B., Martin, E.W., Rhoads, S.N., Reeb, A.N., Nourse, A., Ramirez Montero, D., Ryan, V.H., Rohatgi, R. et al. (2018) A single N-terminal phosphomimic disrupts TDP-43 polymerization, phase separation, and RNA splicing. EMBO J, 37.

23. Yoshizawa, T., Ali, R., Jiou, J., Fung, H.Y.J., Burke, K.A., Kim, S.J., Lin, Y., Peeples, W.B., Saltzberg, D., Soniat, M. et al. (2018) Nuclear Import Receptor Inhibits Phase Separation of FUS through Binding to Multiple Sites. Cell, 173, 693–705 e622.

24. Lin, Y., Protter, D.S., Rosen, M.K. and Parker, R. (2015) Formation and Maturation of Phase-Separated Liquid Droplets by RNA-Binding Proteins. Mol Cell, 60, 208–219.

25. Jean-Philippe, J., Paz, S. and Caputi, M. (2013) hnRNP A1: the Swiss army knife of gene expression. Int J Mol Sci, 14, 18999–19024.

26. Molliex, A., Temirov, J., Lee, J., Coughlin, M., Kanagaraj, A.P., Kim, H.J., Mittag, T. and Taylor, J.P. (2015) Phase separation by low complexity domains promotes stress granule assembly and drives pathological fibrillization. Cell, 163, 123–133.

27. Kim, H.J., Kim, N.C., Wang, Y.D., Scarborough, E.A., Moore, J., Diaz, Z., MacLea, K.S., Freibaum, B., Li, S., Molliex, A. et al. (2013) Mutations in prion-like domains in hnRNPA2B1 and hnRNPA1 cause multisystem proteinopathy and ALS. Nature, 495, 467–473.

28. Morgan, C.E., Meagher, J.L., Levengood, J.D., Delproposto, J., Rollins, C., Stuckey, J.A. and Tolbert, B.S. (2015) The First Crystal Structure of the UP1 Domain of hnRNP A1 Bound to RNA Reveals a New Look for an Old RNA Binding Protein. J Mol Biol, 427, 3241–3257.

29. Pancsa, R., Zsolyomi, F. and Tompa, P. (2018) Co-Evolution of Intrinsically Disordered Proteins with Folded Partners Witnessed by Evolutionary Couplings. Int J Mol Sci, 19.

30. Milkovic, N.M. and Mittag, T. (2020) In Skriver, B. K. a. K. (ed.), Intrinsically Disordered Proteins – Methods and Protocols. Springer Nature, Switzerland, Vol. 2142.

31. Muschol, M. and Rosenberger, F. (1997) Liquid-liquid phase separation in supersaturated lysozyme solutions and associated precipitate formation/crystallization. J Chem Phys, 107, 1953–1962.

32. Stanley, H.E. (1971). Oxford University Press, New York.

33. Sengers, J.V. (1982) Phase Transitions Cargèse 1980. Universality of critical phenomena in classical fluids. 1 ed. Plenum Press, New York.

34. Kirby, N., Cowieson, N., Hawley, A.M., Mudie, S.T., McGillivray, D.J., Kusel, M., Samardzic-Boban, V. and Ryan, T.M. (2016) Improved radiation dose efficiency in solution SAXS using a sheath flow sample environment. Acta Crystallogr D Struct Biol, 72, 1254–1266.

35. Martin, E.W., Hopkins, J.B. and Mittag, T. (2020) Small angle x-ray scattering experiments of monodisperse samples close to the solubility limit. arXiv.

36. Hopkins, J.B., Gillilan, R.E. and Skou, S. (2017) BioXTAS RAW: improvements to a free opensource program for small-angle X-ray scattering data reduction and analysis. J Appl Crystallogr, 50, 1545–1553.

37. Zhao, H., Brautigam, C.A., Ghirlando, R. and Schuck, P. (2013) Overview of current methods in sedimentation velocity and sedimentation equilibrium analytical ultracentrifugation. Curr Protoc Protein Sci, Chapter 20, Unit20 12.

38. Schuck, P. (2016), Sedimentation velocity analytical ultracentrifugation. CRC Press, Boca Raton, FL.

39. Schuck, P. (2000) Size-distribution analysis of macromolecules by sedimentation velocity ultracentrifugation and lamm equation modeling. Biophys J, 78, 1606–1619.

40. Zhao, H., Ghirlando, R., Alfonso, C., Arisaka, F., Attali, I., Bain, D.L., Bakhtina, M.M., Becker, D.F., Bedwell, G.J., Bekdemir, A. et al. (2015) A multilaboratory comparison of calibration accuracy and the performance of external references in analytical ultracentrifugation. PLoS One, 10, e0126420.

41. Brown, P.H. and Schuck, P. (2006) Macromolecular size-and-shape distributions by sedimentation velocity analytical ultracentrifugation. Biophys J, 90, 4651–4661.

42. Sali, A. and Blundell, T.L. (1993) Comparative protein modelling by satisfaction of spatial restraints. J Mol Biol, 234, 779–815.

43. Bayer, P., Arndt, A., Metzger, S., Mahajan, R., Melchior, F., Jaenicke, R. and Becker, J. (1998) Structure determination of the small ubiquitin-related modifier SUMO-1. J Mol Biol, 280, 275–286.

44. Shamoo, Y., Krueger, U., Rice, L.M., Williams, K.R. and Steitz, T.A. (1997) Crystal structure of the two RNA binding domains of human hnRNP A1 at 1.75 A resolution. Nat Struct Biol, 4, 215–222.

45. Monticelli, L., Kandasamy, S.K., Periole, X., Larson, R.G., Tieleman, D.P. and Marrink, S.J. (2008) The MARTINI Coarse-Grained Force Field: Extension to Proteins. J Chem Theory Comput, 4, 819–834.

46. Abraham, M.J., Murtola, T., Schulz, R., Páll, S., Smith, J., Hess, B. and Lindahl, E. (2015) GROMACS: High performance molecular simulations through multi-level parallelism from laptops to supercomputers. SoftwareX, 1-2, 19–25.

47. de Jong, D.H., Singh, G., Bennett, W.F., Arnarez, C., Wassenaar, T.A., Schafer, L.V., Periole, X., Tieleman, D.P. and Marrink, S.J. (2013) Improved Parameters for the Martini Coarse-Grained Protein Force Field. J Chem Theory Comput, 9, 687–697.

48. Wassenaar, T.A., Ingolfsson, H.I., Bockmann, R.A., Tieleman, D.P. and Marrink, S.J. (2015) Computational Lipidomics with insane: A Versatile Tool for Generating Custom Membranes for Molecular Simulations. JChem Theory Comput, 11, 2144–2155.

49. Larsen, A.H., Wang, Y., Bottaro, S., Grudinin, S., Arleth, L. and Lindorff-Larsen, K. (2019) Combining molecular dynamics simulations with small-angle X-ray and neutron scattering data to study multi-domain proteins in solution. bioRxiv.

50. Wassenaar, T.A., Pluhackova, K., Bockmann, R.A., Marrink, S.J. and Tieleman, D.P. (2014) Going Backward: A Flexible Geometric Approach to Reverse Transformation from Coarse Grained to Atomistic Models. J Chem Theory Comput, 10, 676–690.

51. Grudinin, S., Garkavenko, M. and Kazennov, A. (2017) Pepsi-SAXS: an adaptive method for rapid and accurate computation of small-angle X-ray scattering profiles. Acta Crystallogr D Struct Biol, 73, 449–464.

52. Bottaro, S., Bengtsen, T. and Lindorff-Larsen, K. (2020) Integrating Molecular Simulation and Experimental Data: A Bayesian/Maximum Entropy Reweighting Approach. Methods Mol Biol, 2112, 219–240.

53. Ryan, V.H., Dignon, G.L., Zerze, G.H., Chabata, C.V., Silva, R., Conicella, A.E., Amaya, J., Burke, K.A., Mittal, J. and Fawzi, N.L. (2018) Mechanistic View of hnRNPA2 Low-Complexity Domain Structure, Interactions, and Phase Separation Altered by Mutation and Arginine Methylation. Mol Cell, 69, 465–479 e467.

54. Maurer, R.W., Sandler, S.I. and Lenhoff, A.M. (2011) Salting-in characteristics of globular proteins. Biophys Chem, 156, 72–78.

55. Orioli, S., Larsen, A.H., Bottaro, S. and Lindorff-Larsen, K. (2020) How to learn from inconsistencies: Integrating molecular simulations with experimental data. Prog Mol Biol Transl Sci, 170, 123–176.

56. Brangwynne, C.P., Tompa, P. and Pappu, R.V. (2015) Polymer physics of intracellular phase transitions. Nature Physics, 11, 899–904.

57. Alberti, S., Saha, S., Woodruff, J.B., Franzmann, T.M., Wang, J. and Hyman, A.A. (2018) A User’s Guide for Phase Separation Assays with Purified Proteins. J Mol Biol, 430, 4806–4820.

58. Schmidt, H.B., Barreau, A. and Rohatgi, R. (2019) Phase separation-deficient TDP43 remains functional in splicing. Nat Commun, 10, 4890.

59. Hofweber, M., Hutten, S., Bourgeois, B., Spreitzer, E., Niedner-Boblenz, A., Schifferer, M., Ruepp, M.D., Simons, M., Niessing, D., Madl, T. et al. (2018) Phase Separation of FUS Is Suppressed by Its Nuclear Import Receptor and Arginine Methylation. Cell, 173, 706–719 e713.

60. Guo, L., Kim, H.J., Wang, H., Monaghan, J., Freyermuth, F., Sung, J.C., O’Donovan, K., Fare, C.M., Diaz, Z., Singh, N. et al. (2018) Nuclear-Import Receptors Reverse Aberrant Phase Transitions of RNA-Binding Proteins with Prion-like Domains. Cell, 173, 677–692 e620.

61. Qamar, S., Wang, G., Randle, S.J., Ruggeri, F.S., Varela, J.A., Lin, J.Q., Phillips, E.C., Miyashita, A., Williams, D., Strohl, F. et al. (2018) FUS Phase Separation Is Modulated by a Molecular Chaperone and Methylation of Arginine Cation-pi Interactions. Cell, 173, 720–734 e715.

62. Dahal, Y.R. and Schmit, J.D. (2018) Ion Specificity and Nonmonotonic Protein Solubility from Salt Entropy. Biophys J, 114, 76–87.

63. Yang, P., Mathieu, C., Kolaitis, R.M., Zhang, P., Messing, J., Yurtsever, U., Yang, Z., Wu, J., Li, Y., Pan, Q. et al. (2020) G3BP1 Is a Tunable Switch that Triggers Phase Separation to Assemble Stress Granules. Cell, 181, 325–345 e328.

64. Guillen-Boixet, J., Kopach, A., Holehouse, A.S., Wittmann, S., Jahnel, M., Schlussler, R., Kim, K., Trussina, I., Wang, J., Mateju, D. et al. (2020) RNA-Induced Conformational Switching and Clustering of G3BP Drive Stress Granule Assembly by Condensation. Cell, 181, 346–361 e317.

65. Mitrea, D.M., Cika, J.A., Stanley, C.B., Nourse, A., Onuchic, P.L., Banerjee, P.R., Phillips, A.H., Park, C.G., Deniz, A.A. and Kriwacki, R.W. (2018) Self-interaction of NPM1 modulates multiple mechanisms of liquid-liquid phase separation. Nat Commun, 9, 842.

66. Rayman, J.B., Karl, K.A. and Kandel, E.R. (2018) TIA-1 Self-Multimerization, Phase Separation, and Recruitment into Stress Granules Are Dynamically Regulated by Zn(2). Cell Rep, 22, 59–71.

