## Supplementary Data for "Interplay of folded domains and the disordered low-complexity domain in mediating hnRNPA1 phase separation"

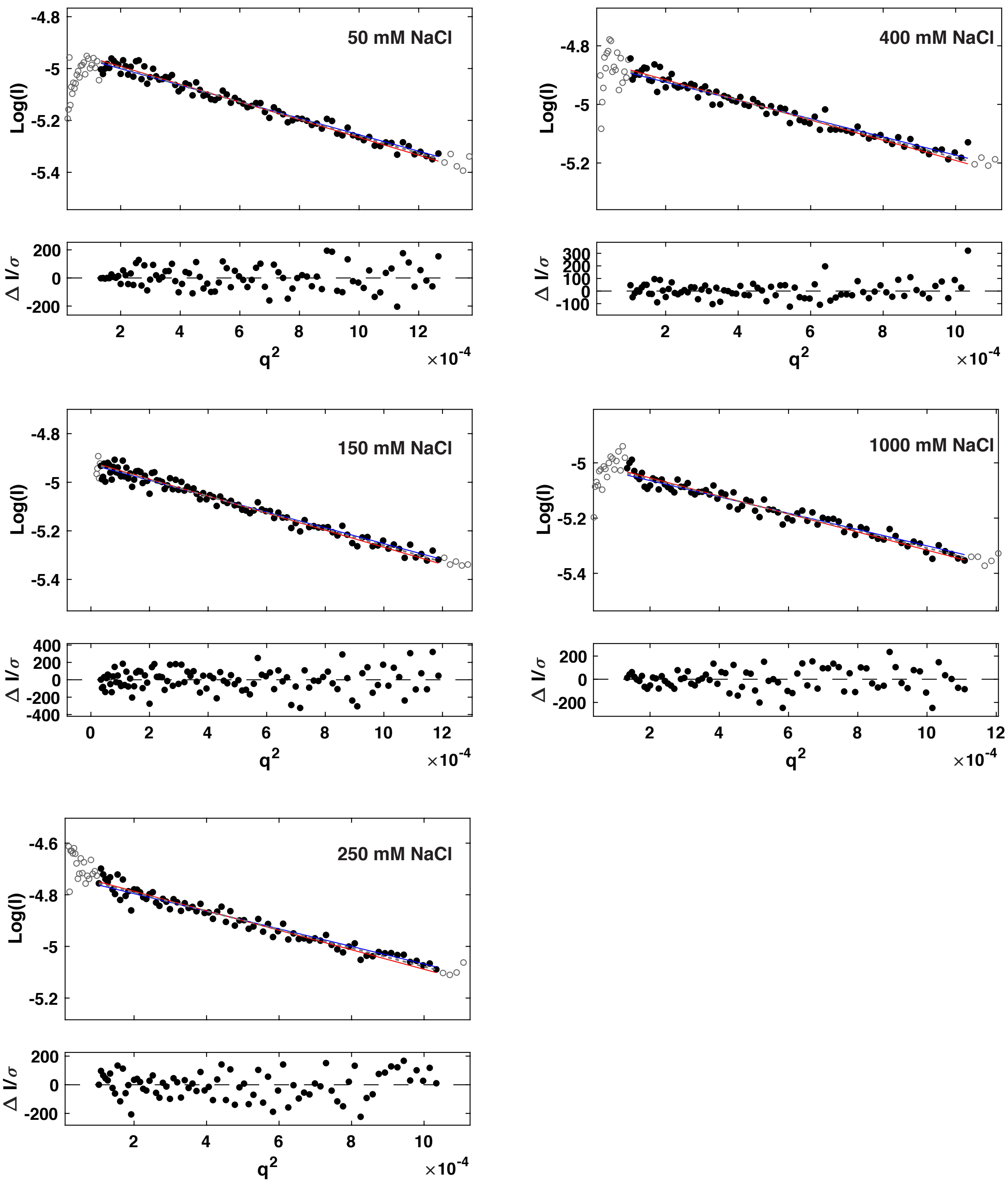


**Figure S1: Guinier analysis of SAXS data on samples of hnRNPA1* shown in Fig. 3.**
Guinier fits (dashed lines) to raw data (filled circles); raw data outside the fitted q-range are shown as open circles. Steepest and most shallow possible linear fits to the data are shown as red and blue lines, respectively. The lower panels show the residuals to the linear fit.

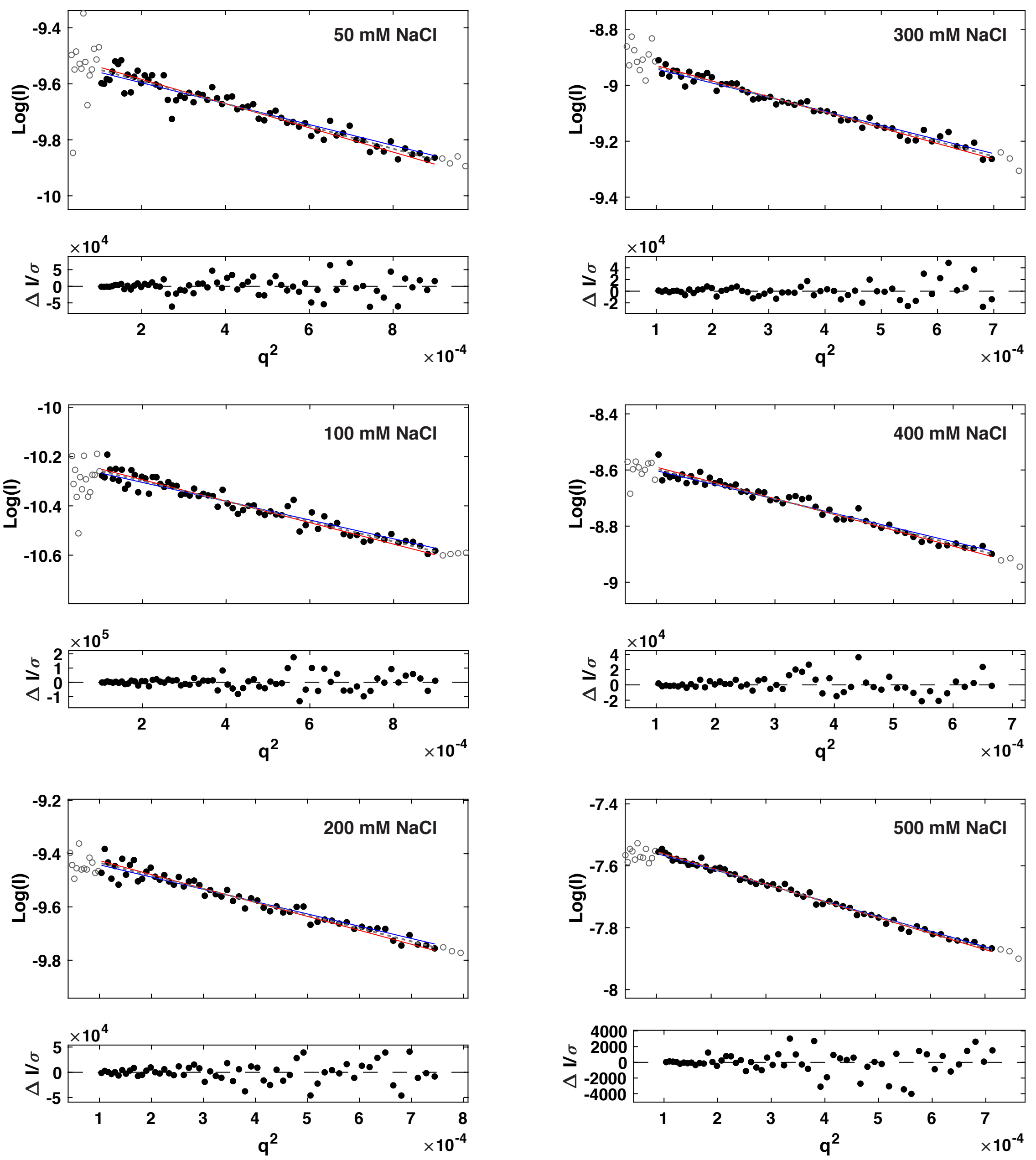


**Figure S2: Guinier analysis of SAXS data on samples of hSUMO-hnRNPA1* shown in Fig. 4.**
Guinier fits (dashed lines) to raw data (filled circles); raw data outside the fitted q-range are shown as open circles. Steepest and most shallow possible linear fits to the data are shown as red and blue lines, respectively. The lower panels show the residuals to the linear fit.

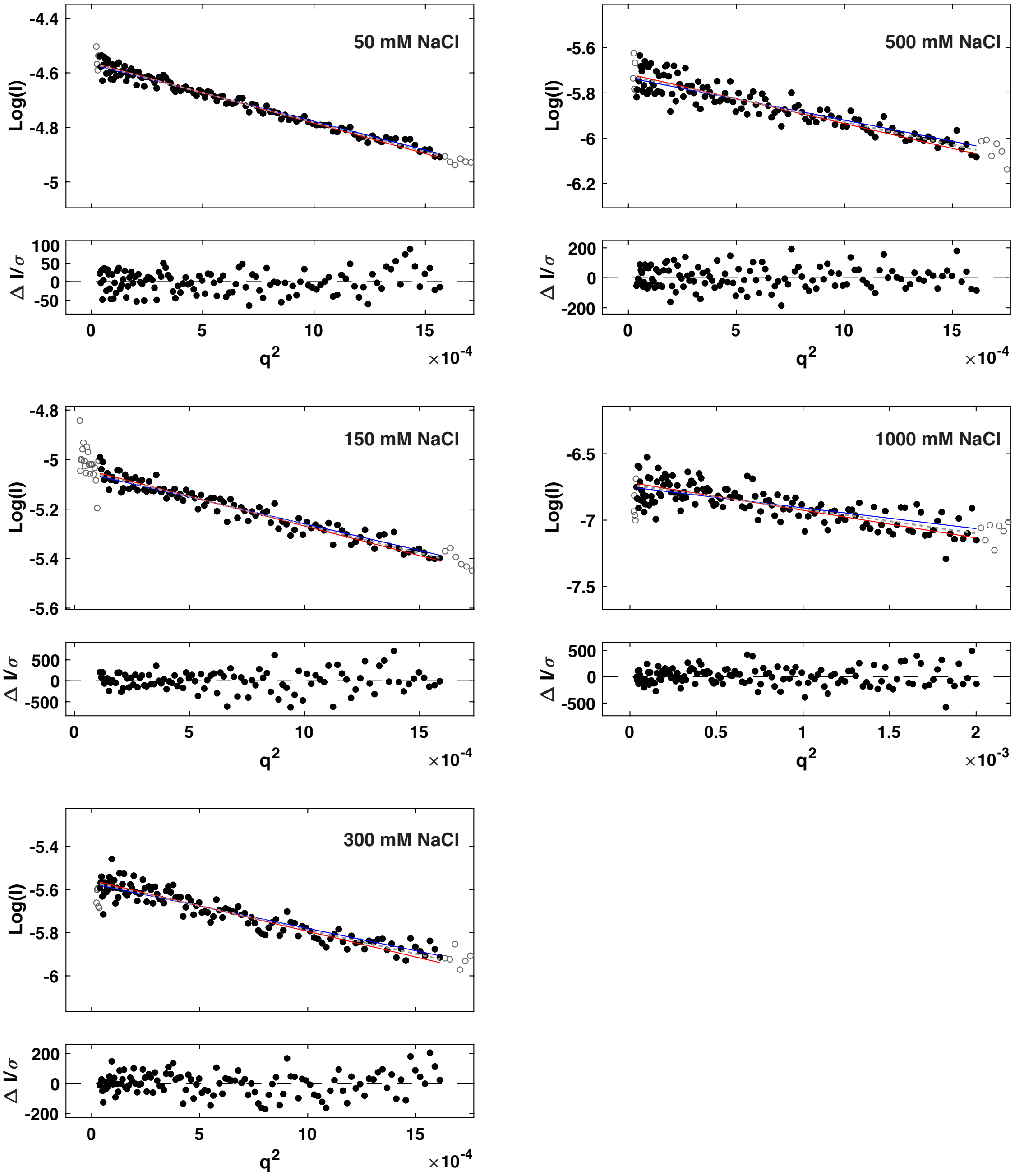


**Figure S3: Guinier analysis of SAXS data on samples of LCD* shown in Fig. 5.**
Guinier fits (dashed lines) to raw data (filled circles); raw data outside the fitted q-range are shown as open circles. Steepest and most shallow possible linear fits to the data are shown as red and blue lines, respectively. The lower panels show the residuals to the linear fit.

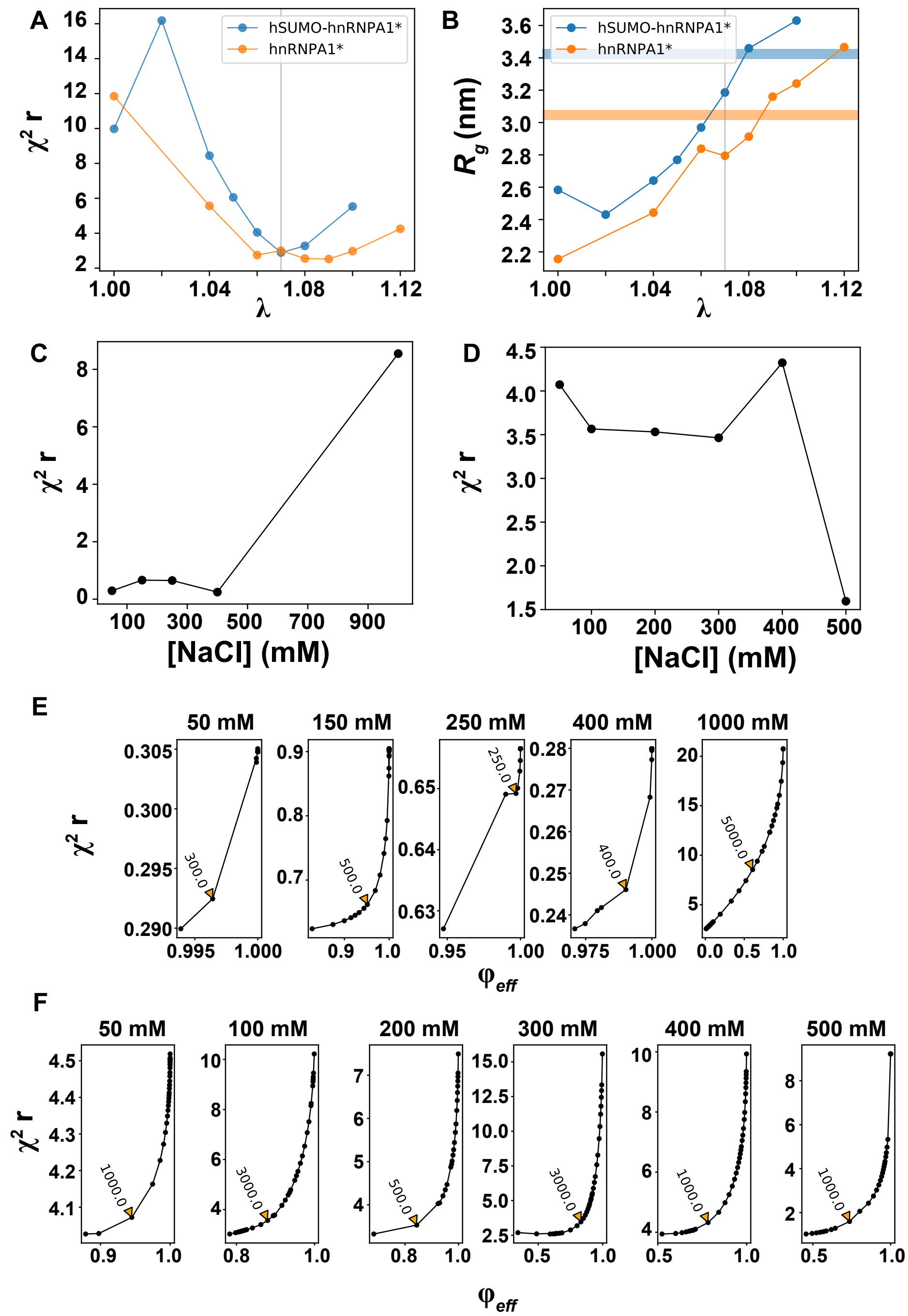


**Figure S4: Tuning protein-water interactions and extent of reweighting of coarse-grained MD simulations.**
**(A)** The protein-water interaction parameter λ was changed progressively in simulations of hnRNPA1* and hSUMO-hnRNPA1* at 150 mM and 100 mM NaCl, respectively, and the fit of the resulting ensembles to the experimental SAXS data determined. λ of 1.07 (i.e., the value that gave the best agreement with experiment; grey bars) was chosen for further simulations.
**(B)** The $R_g$ values from simulation and experiment are similar at λ of 1.07. The orange and blue bars show the experimental $R_g$ values for hnRNPA1* and hSUMO-hnRNPA1*, respectively.
**(C)** Fit of simulations to experimental SAXS data after reweighting for hnRNPA1*.
**(D)** Fit of simulations to experimental SAXS data after reweighting for hSUMO-hnRNPA1*.
**(E)** The extent of reweighting optimized a balance between the fit to the experimental SAXS data and usage of a large fraction of the trajectory for hnRNPA1*.
**(F)** The extent of reweighting optimized a balance between the fit to the experimental SAXS data and usage of a large fraction of the trajectory for hSUMO-hnRNPA1*.

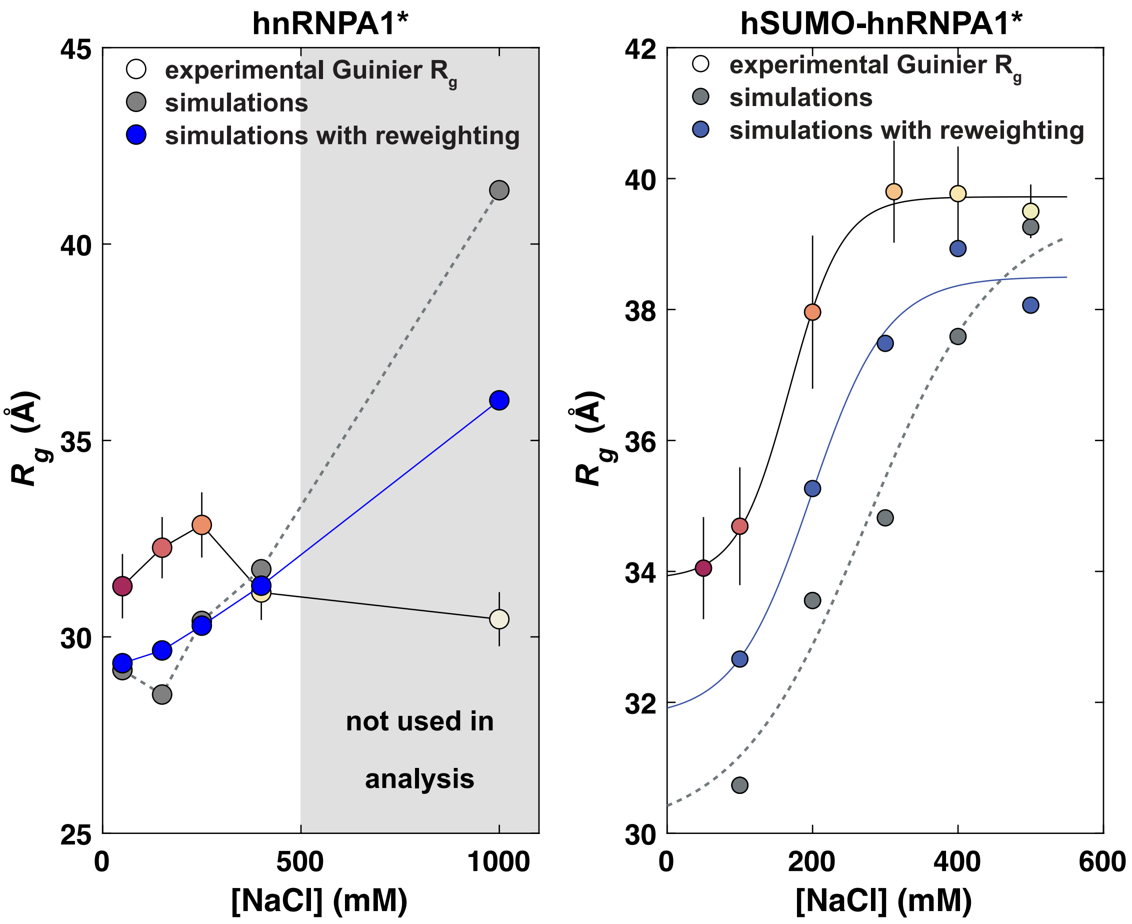


**Figure S5: Comparison of $R_\mathrm{g}$ values from Guinier analysis and MD simulations without and with reweighting for hnRNPA1* and hSUMO-hnRNPA1*.**

$R_\mathrm{g}$ values from Guinier analysis are based on a small portion of the experimental data in the Guinier q-range (see also Fig. S1 and S2). When experimental data is interpreted via MD simulations, the whole q-range is used for reweighting (see also Fig. 3 and 4). The data for hSUMO-hnRNPA1* was fit to logistic functions that indicate minimum dimensions at low NaCl concentration, a maximum extension at high NaCl concentration and a transition with a midpoint at ~175 mM NaCl, suggesting screening of interactions between oppositely charged regions of hSUMO-hnRNPA1*. Simulations recapitulate the trend and after reweighting show close agreement.

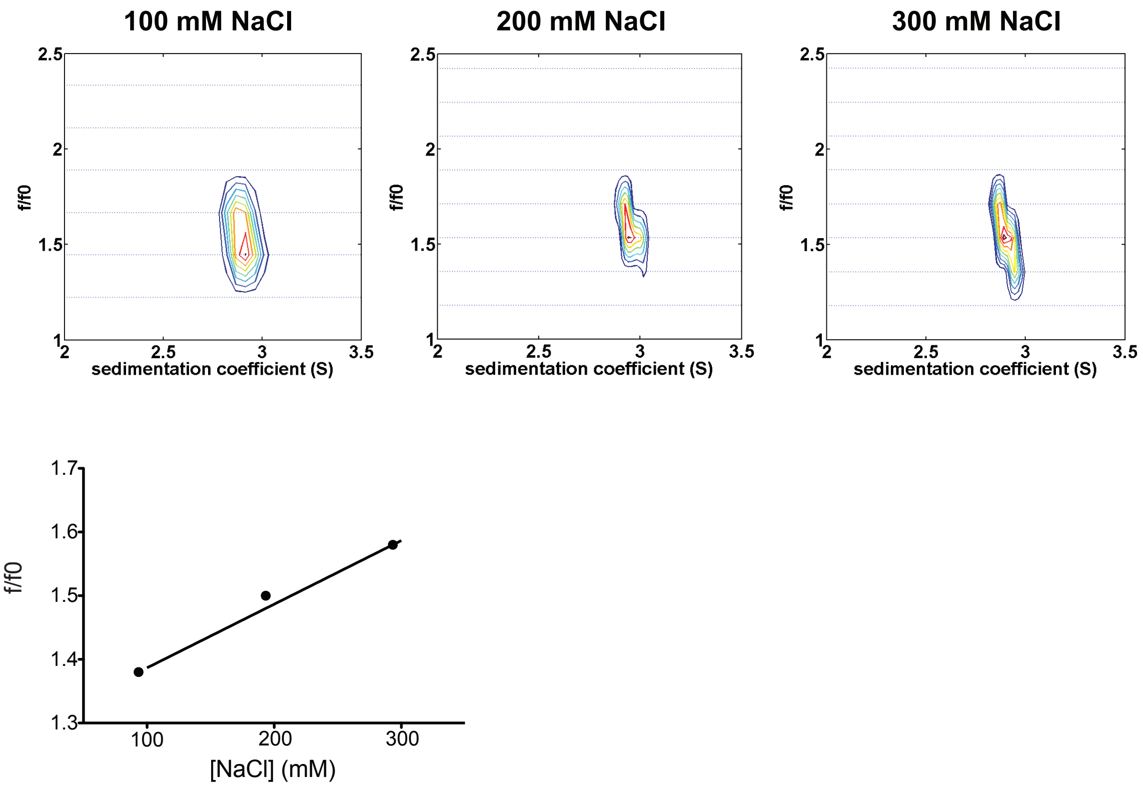


**Figure S6: Sedimentation velocity analysis of hnRNPA1 as a function of NaCl concentration.**
The sedimentation velocity profiles (fringe displacement) were fitted to a two-dimensional size-and-shape distribution model and shown as contour plots (heat maps) of $c(s,f/f_0)$ with increasing color temperature indicating higher fringes/S values. Velocity data were acquired at a rotor speed of 50,000 rpm at 20 °C in 50 mM HEPES pH 7.5, 5 mM DTT and 100, or 200 or 300 mM NaCl, buffer at 20 °C. Rayleigh interference optical data were collected.
